# Human coronavirus HKU1 recognition of the TMPRSS2 host receptor

**DOI:** 10.1101/2024.01.09.574565

**Authors:** Matthew McCallum, Young-Jun Park, Cameron Stewart, Kaitlin R. Sprouse, Jack Brown, M. Alejandra Tortorici, Cecily Gibson, Emily Wong, Margareta Ieven, Amalio Telenti, David Veesler

## Abstract

The human coronavirus HKU1 spike (S) glycoprotein engages host cell surface sialoglycans and transmembrane protease serine 2 (TMPRSS2) to initiate infection. The molecular basis of HKU1 binding to TMPRSS2 and determinants of host receptor tropism remain elusive. Here, we designed an active human TMPRSS2 construct enabling high-yield recombinant production in human cells of this key therapeutic target. We determined a cryo-electron microscopy structure of the HKU1 RBD bound to human TMPRSS2 providing a blueprint of the interactions supporting viral entry and explaining the specificity for TMPRSS2 among human type 2 transmembrane serine proteases. We found that human, rat, hamster and camel TMPRSS2 promote HKU1 S-mediated entry into cells and identified key residues governing host receptor usage. Our data show that serum antibodies targeting the HKU1 RBD TMPRSS2 binding-site are key for neutralization and that HKU1 uses conformational masking and glycan shielding to balance immune evasion and receptor engagement.

---

HKU1 is one of the four β-coronaviruses known to circulate in humans seasonally. It was initially discovered in 2005 in nasopharyngeal aspirates obtained from a 71-year-old man with pneumonia in Hong Kong^1^. Retrospective investigations identified HKU1 in nasopharyngeal aspirates collected between 1995 and 2002 from symptomatic individuals on several continents^1–4^ and more HKU1 strains were sequenced and isolated thereafter^5, 6^. Although HKU1 is largely considered to be a ‘common cold’ coronavirus, mainly causing mild respiratory infections, it also causes severe illness, particularly (but not only) for children, the elderly and immunocompromised individuals^7–9^. Furthermore, its high seroprevalence in multiple independent studies indicates that HKU1 continuously circulates in the human population and is therefore endemic^8^.

Coronavirus infections are initiated by interaction of the trimeric spike (S) glycoprotein with a host cell receptor leading to fusion of the viral and host membrane and genome delivery^10^. To date, only a finite number of proteinaceous receptors have been identified as promoting coronavirus infection and all of them are host transmembrane proteases: angiotensin-converting enzyme 2 (ACE2) for SARS-CoV-1^11^, SARS-CoV-2^12–14^, NL63^15^ and a few divergent bat-borne merbecoviruses^16^; aminopeptidase N (APN) for 229E^17^, CcoV-HuPn-2018^18^, PDCoV^19^ and TGEV/PEDV^20^; dipeptidyl-peptidase 4 (DPP4) for MERS-CoV^21^ and HKU4-related merbecoviruses^22–24^; and carcinoembryonic antigen-related cell adhesion molecule 1a for MHV A59^25^. Furthermore, OC43 and HKU1 recognize 9-O-acetylated sialosides^26–28^ present at the host cell surface and a recent study showed that transmembrane protease serine 2 (TMPRSS2) is an entry receptor for HKU1^29^, which is likely engaged upon S conformational changes triggered by α2,8-linked 9-O-acetylated disialosides binding^30^. The molecular basis of HKU1 recognition of the human TMPRSS2 receptor along with the determinants of host receptor tropism, however, remain elusive.

TMPRSS2 is part of the type 2 transmembrane serine protease family that encompasses several cell surface-expressed trypsin-like serine proteases participating in epithelial homeostasis and TMPRSS2 genomic rearrangements are common in prostate cancer^31^. Furthermore, TMPRSS2 has been shown to proteolytically activate the S glycoprotein of many coronaviruses, including SARS-CoV-2^32, 33^, SARS-CoV-1^34, 35^, MERS-CoV^36, 37^, 229E^38^, as well as influenza virus hemagglutinins^39, 40^. As a result, TMPRSS2 inhibitors have been proposed as candidates for preventing or treating these viral infections^41–43^. Recombinant production of TMPRSS2 and related type 2 transmembrane serine proteases has proven exceptionally challenging, hindering studies of these important therapeutic targets.

Here, we designed a human TMPRSS2 glycoprotein construct enabling high-yield recombinant production of the active mature enzyme ectodomain in human cells, which will expedite biochemical studies and drug screening globally. We determined a cryo-electron microscopy (cryoEM) structure of the human TMPRSS2 ectodomain bound to the HKU1 S receptor-binding domain (RBD), providing a blueprint of the interactions mediating receptor engagement, specificity and host receptor tropism. Analysis of HKU1 infection-elicited serum antibodies in humans reveal that the TMPRSS2-binding site is a key site of vulnerability for HKU1 that becomes exposed upon sialoglycan-mediated RBD opening. Our findings reveal how this human coronavirus gains access to target host cells and balances receptor recognition and immune evasion, informing vaccine design.

## Design of an active human TMPRSS2 construct

Given the therapeutic relevance of human TMPRSS2, we sought to design a strategy enabling recombinant production of native enzyme from human cells. Although a recent study reported producing the active TMPRSS2 ectodomain (dasTMPRSS2) using Sf9 cells^44^, insect cell expression is inherently time-consuming and does not faithfully recapitulate the chemical composition of protein glycosylations. Furthermore, the replacement of the native TMPRSS2 autoactivation cleavage site with an enterokinase cleavage sequence in dasTMPRSS2 removes the endogeneous N249 glycosylation motif and also requires an additional proteolytic activation step, adding to the laborious nature of this purification.

To overcome these limitations, we designed an ectodomain construct enabling high-yield and rapid recombinant production of active human TMPRSS2. We first observed that we could not detect dasTMPRSS2 expression in Expi293 human cells even after introducing the S441A residue mutation to disable proteolytic activity and potential cytotoxicity during expression (**Figure 1A-B, Figure S1 and Table S1**). Human TMPRSS2 harbors an unpaired and solvent-exposed cysteine at position 379 in the protease domain, which otherwise participates in a disulfide bond with the residue equivalent to T447 in most related type 2 transmembrane serine proteases^44^. We therefore introduced the T447C residue substitution to allow formation of a C379-C447 disulfide bond, which was sufficient to detect TMPRSS2 expression, albeit with low yield (**Figure 1A-B, Figure S1 and Table S1**). Fusion of an enterokinase-cleavable N-terminal SUMO tag increased production yield approximately 2-fold, relative to the construct lacking the SUMO fusion (**Figure 1A-B, Figure S1 and Table S1**). Finally, reintroduction of two native serine residues, corresponding to positions 250 and 251 of the human TMPRSS2 sequence, restored the N-linked glycan at position N249 and improved production yield 8-fold compared to the glycan knockout construct (**Figure 1A-B, Figure S1 and Table S1**).

**Figure 1.**
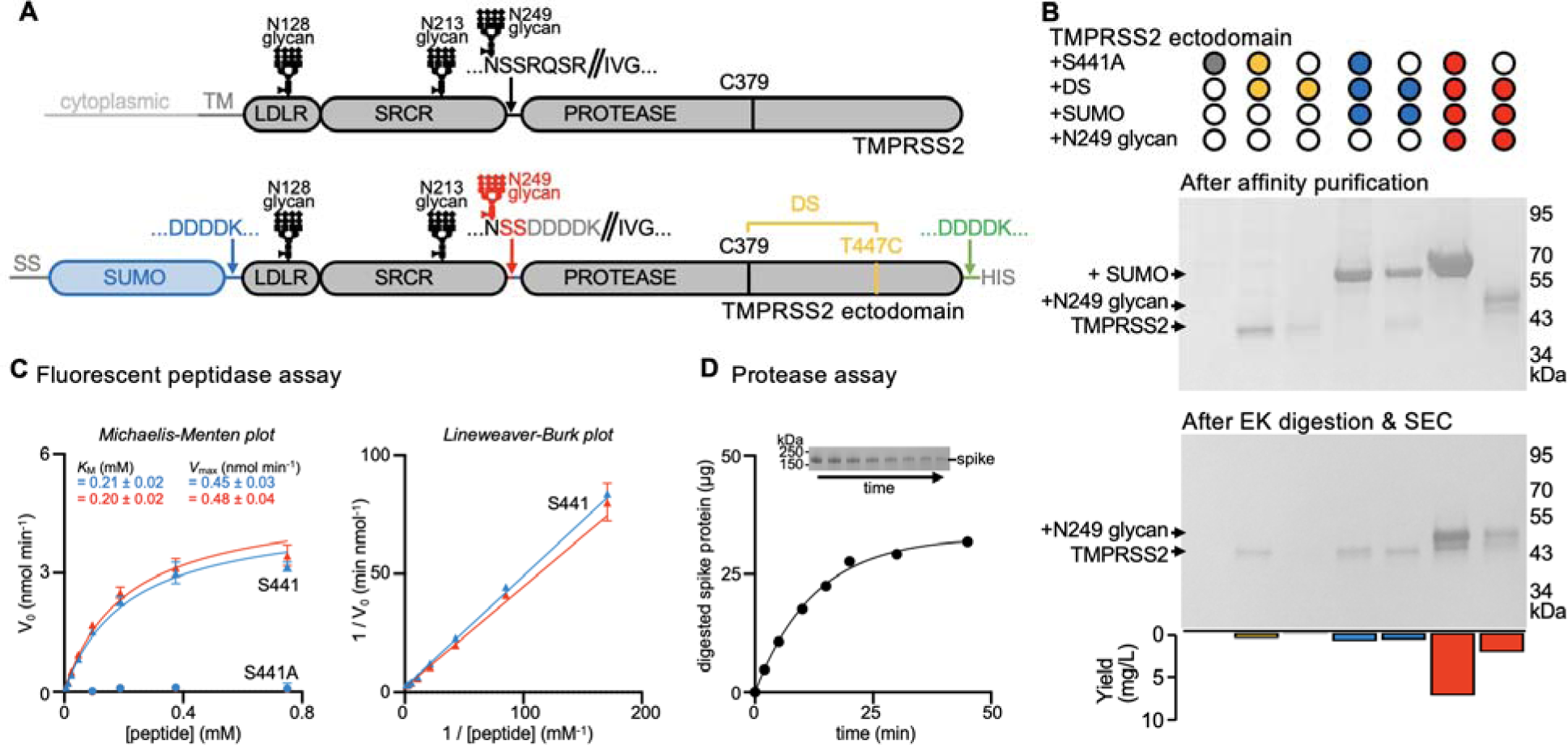
Design and functional characterization of an enzymatically active human TMPRSS2 construct. (A), Schematic representation of the domain architecture of human TMPRSS2 and of the ectodomain construct designed for mammalian expression. The sequences of the constructs tested can be found in **Table S1**. Designed features are a C379-T447C disulfide bond (DS, yellow), an enterokinase-cleavable N-terminal SUMO fusion (blue), and reintroduction of the N249 glycosylation motif (red) relative to a previously described insect cell expression construct (dasTMPRSS2)^44^. LDLR: low-density lipoprotein receptor domain; SRCR: Scavenger Receptor Cysteine-Rich domain; PROTEASE: trypsin-like serine protease domain. SS: signal sequence. DDDDK: enterokinase cleavage sequence. (B) Non-reducing SDS-PAGE analysis of purified TMPRSS2 constructs with or without S441A, the C379-T447C disulfide bond (DS), an N-terminal SUMO fusion, and the native glycan at position N249. Proteins were expressed and affinity purified in parallel, and then normalized to a standard volume to enable comparisons of relative yields and purity after affinity purification (top) and after enterokinase (EK) digestion and SEC purification (middle). The final yield of purified TMPRSS2 per liter of Expi293 expression medium is indicated as a bar graph (bottom). (C) Michaelis–Menten plot (left) and Lineweaver-Burke plot (right) of initial reaction velocities with various concentrations of fluorescent Boc-QAR-AMC peptide substrate, at 22 °C, in the presence of 6.8 nM of the TMPRSS2 ectodomain harboring either the DS and SUMO (blue) or the DS, SUMO, and N249 glycan (red) modifications. *K_M_* and *V_max_* were determined through fitting using GraphPad Prism. Data are shown as the geometric mean and standard deviation of three technical replicates. (D) Proteolytic processing of 40 µg of SARS-CoV-2 S 2P (S2P) by 0.5 µM TMPRSS2 over 45 min at 37°C, visualized by reducing SDS-PAGE analysis (inset) and measured by densitometry. The inset on top shows the decreasing intensity of the S2P band over time used for these calculations.

Reintroduction of the S441 catalytic residue reduced yields ∼3 fold relative to S441A-harboring TMPRSS2 (**Figure 1A-B, Figure S1 and Table S1**). We noted that the TMPRSS2 construct harboring the wildtype S441 residue, T447C, the N-terminal SUMO fusion, and the native N249 glycan autocatalytically processed the enterokinase cleavage sequences during expression, leading to zymogen activation and SUMO cleavage (**Figure 1A-B, Figure S1 and Table S1**). Consistent with this interpretation, the catalytically inactive S441A-harboring TMPRSS2 mutant remains uncleaved. Overall, our designed constructs streamline TMPRSS2 production yielding 7 mg of purified S441A TMPRSS2 or 2 mg of self-activating S441 TMPRSS2 per liter of human Expi293 cells, 4 days after transfection, which will accelerate studies of this key therapeutic target.

Purified TMPRSS2 is highly active in a fluorescent peptidase assay, using the fluorogenic Boc-Gln-Ala-Arg-7-aminomethylcoumarine (Boc-QAR-AMC) peptide substrate, and its activity can be modeled with Michaelis-Menten kinetics (**Figure 1C and Figure S1**). We determined a *K_M_* of 200 µM and V_max_ of 0.5 nmol/min for Boc-QAR-AMC, similar to dasTMPRSS2 purified from insect cells^44^. Given that reintroduction of the N249 glycan did not significantly alter TMPRSS2 enzyme kinetics, the observed autocatalytic cleavage likely resulted from a high TMPRSS2 concentration during production rather than enhanced activity (**Figure 1C and Figure S1**). TMPRSS2 is also active using the SARS-CoV-2 S ectodomain trimer as a substrate (**Figure 1D**).

## Structure of the TMPRSS2-bound HKU1 RBD

To understand the molecular basis of HKU1 engagement of its host receptor, we characterized an HKU1 isolate N1 (genotype A) RBD in complex with the human TMPRSS2 ectodomain harboring the S441A substitution of the catalytic nucleophile. We used cryoEM to determine a structure of this 85kDa complex at 2.9 Å resolution revealing that the TMPRSS2 catalytic domain is engaged by the HKU1 receptor-binding motif (RBM) which is inserted in between two β-strands of the RBD core β-sheet (**Figure 2A-D, Figure S2 and Table S2**). No major conformational changes occur upon binding besides small scale structural rearrangements of the HKU1 RBM mostly centered on the β-hairpin spanning residues 505-517, as compared to the isolated RBD structure^45^ (**Figure S3**).

**Figure 2.**
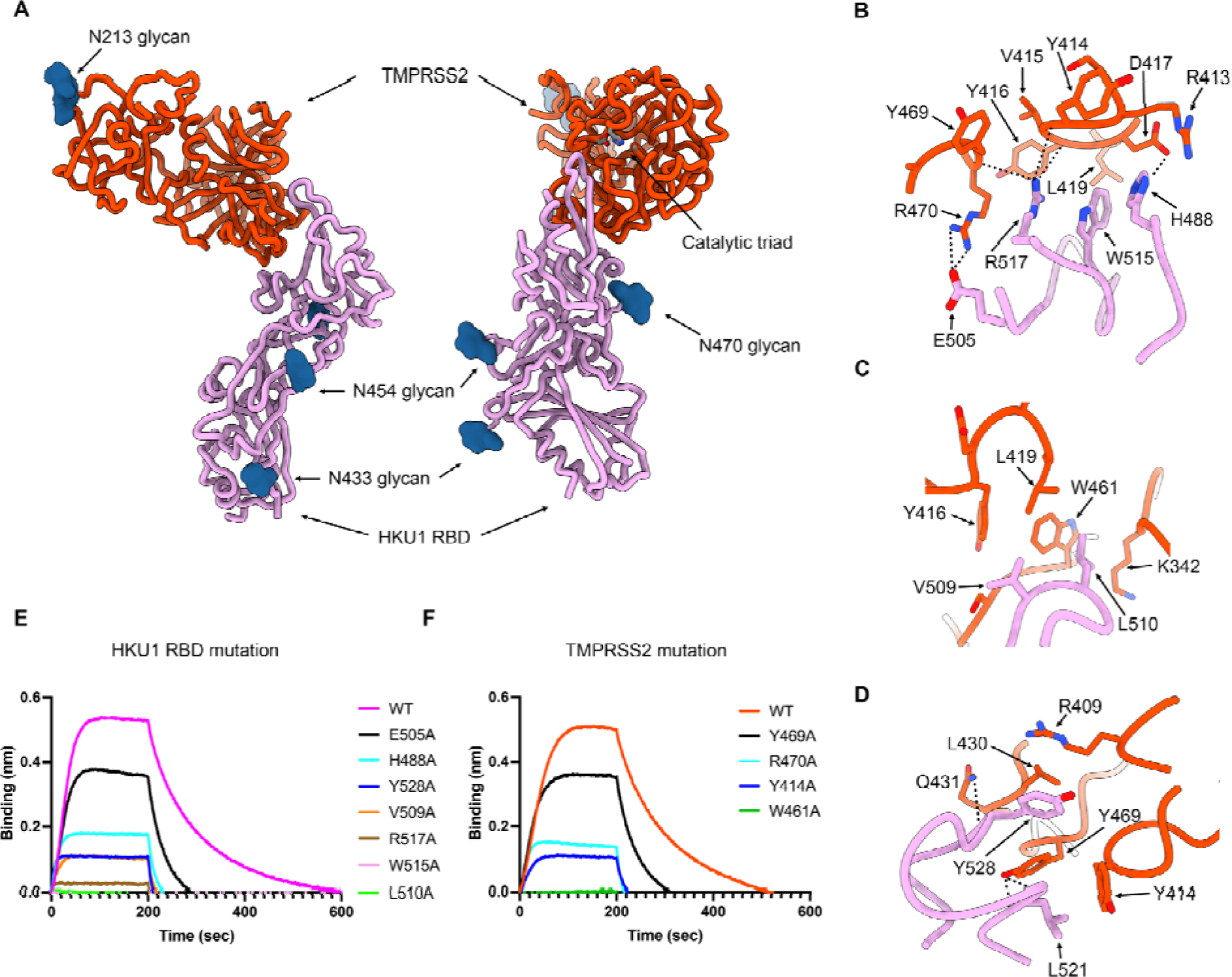
Molecular basis of human TMPRSS2 recognition by the HKU1 RBD. (A) Ribbon diagrams in two orthogonal orientations of the cryoEM structure of the HKU1 RBD (purple) bound to the human TMPRSS2 ectodomain (orange) at 2.9Å resolution. The TMPRSS2 catalytic triad residues (H296, D345 and S441) are shown as sticks colored by heteroatom and N-linked glycans are rendered as blue spheres. (B-D) Zoomed-in views of the interface highlighting key interactions between the HKU1 RBD and the human TMPRSS2 peptidase domain. Select electrostatic interactions are shown as black dotted lines. (E) Binding of S441A TMPRSS2 to the wildtype (WT) isolate N1 and to the H488A, E505A, V509A, L510A, W515A, R517A and Y528A HKU1 RBD interface mutants immobilized on biolayer interferometry SA biosensors. (F) Binding of the wildtype (WT) and the Y414A, W461A, Y469A and R470A human TMPRSS2 (active S441) interface mutants to the wildtype isolate N1 HKU1 RBD immobilized on biolayer interferometry SA biosensors.

Receptor engagement leads to burial of surfaces of approximately 800 Å^2^ from each of the two binding partners using electrostatic interactions and surface complementarity. The HKU1 RBM comprises residues 488, 505, 507-512, 515, 517-522, 527-532, and 554 to recognize TMPRSS2 residues 340-342, 409, 412-417, 419, 430-431, 433, 461-463 and 468-470 forming loops protruding from one of the two β-barrels of the protease domain (**Figure 2A**). Key interactions include (i) the H488_HKU1_ imidazole side chain salt-bridged to the D417_TMPRSS2_ side chain carboxylate (**Figure 2B**); (ii) the E505_HKU1_ side chain carboxylate forming a salt bridge with the R470_TMPRSS2_ side chain guanidinium (**Figure 2B**); (iii) the V509_HKU1_ side chain establishing van der Waals contacts with the Y416_TMPRSS2_, L419_TMPRSS2_ and W461_TMPRSS2_ side chains (**Figure 2C**); (iv) the L510_HKU1_ side chain inserting in a surface-exposed cleft and contacting the aliphatic part of the K342_TMPRSS2_ side chain and the W461_TMPRSS2_ indol ring (**Figure 2C**); (v) the W515_HKU1_ side chain forming van der Waals contacts with Y416_TMPRSS2_, D417_TMPRSS2_ and L419_TMPRSS2_ (**Figure 2A**); (vi) the R517_HKU1_ side chain electrostatically interacting with the Y414_TMPRSS2_, V415_TMPRSS2_ and Y469_TMPRSS2_ backbone carbonyls (**Figure 2A**); (vii) the L521_HKU1_ side chain which is sandwiched between the Y414_TMPRSS2_ and Y469_TMPRSS2_ side chains and the L521 backbone amide and carbonyl that are hydrogen-bonded to the Y469_TMPRSS2_ side chain phenol (**Figure 2D**); and (viii) the Y528_HKU1_ side chain interacting with the R409_TMPRSS2_, L430_TMPRSS2_ and Y469_TMPRSS2_ side chains and the Y528_HKU1_ backbone carbonyl which is hydrogen-bonded to the Q431_TMPRSS2_ amide side chain (**Figure 2D**). As none of the HKU1 S or TMPRSS2 N-linked glycans contribute to the binding interface, the previously reported HKU1 S pseudovirus sensitivity to neuraminidase treatment of target cells^29^ likely resulted from removal of sialoglycan receptors, thereby dampening entry.

## Validation of the TMPRSS2-bound HKU1 RBD cryoEM structure

To functionally validate the role of key interacting residues identified in our cryoEM structure, we evaluated binding of catalytically active or inactive TMPRSS2 to seven isolate N1 HKU1 RBD point mutants harboring an alanine substitution at the receptor-binding interface. The HKU1 RBD L510A and W515A mutations had the strongest effect among the mutants tested, completely preventing binding (**Figure 2E**). R517A had the next strongest effect, severely abrogating recognition, followed by V509A, Y528A, H488A and E505A (**Figure 2E**). The deleterious impact of the HKU1 RBD W515A and R517A mutations on TMPRSS2 binding concur with prior biochemical data^29, 45^, underscoring the key role of these two viral residues. Furthermore, we found that the TMPRSS2 W461A, Y414A, R470A and Y469A interface mutants had no or reduced binding to the HKU1 RBD (**Figure 2F**). These data demonstrate that the HKU1 S and TMPRSS2 residues identified as main contributors to binding in our cryoEM structure are mediating attachment of the RBD to its host receptor and that mutations of these residues to alanine is deleterious for TMPRSS2 binding.

## HKU1 RBD binding inhibits TMPRSS2 activity

Upon binding, the HKU1 RBM β-hairpin (comprising residues 505-517) is positioned in close proximity to the active site albeit without contacting the catalytic triad residues (H296, D345 and S441) (**Figure 2A**). Superimposing the HKU1 RBD-bound TMPRSS2 structure with that of the KQLR chloromethylketone-bound human hepsin^46^ indicates that the HKU1 RBM would compete with binding of this substrate to the TMPRSS2 active site through steric hindrance, assuming an identical binding mode of the ligand (**Figure 3A**). To assess a possible impact on TMPRSS2 enzymatic activity, we quantified proteolytic processing of the Boc-QAR-AMC peptide substrate in the presence of various concentrations of the HKU1 RBD. TMPRSS2 proteolytic activity was inhibited by the HKU1 RBD in a dose-dependent manner with an inhibition constant K*_I_* of 38 +/- 5 nM (**Figure 3B-D and Figure S4**). Furthermore, the seven HKU1 RBD alanine point mutants at the receptor-binding interface had dampened inhibitory effect on TMPRSS2 activity (**Figure 3B and Figure S4**), concurring with the binding reductions observed by biolayer interferometry (**Figure 2E**). The W515A HKU1 RBD mildly increased TMPRSS2 peptidase activity, as was the case for bovine serum albumin (BSA), suggesting non-specific TMPRSS2 stabilization. Michaelis-Menten kinetics analysis of TMPRSS2 activity in the presence of various concentrations of the HKU1 RBD suggested a mechanism of action through competitive inhibition (**Figure 3C-D**), in line with partial occluding of the active site observed structurally. Furthermore, the HKU1 RBD inhibited cleavage of the purified SARS-CoV-2 S 2P trimer in a dose-dependent manner, as evaluated by SDS-PAGE (**Figure 2F**). These results underscore the lack of requirement for TMPRSS2 enzymatic activity to promote HKU1 entry, as the R255Q and the S441A inactive mutants were shown to support HKU1 S-mediated pseudovirus entry with comparable levels to wildtype TMPRSS2^29^.

**Figure 3.**
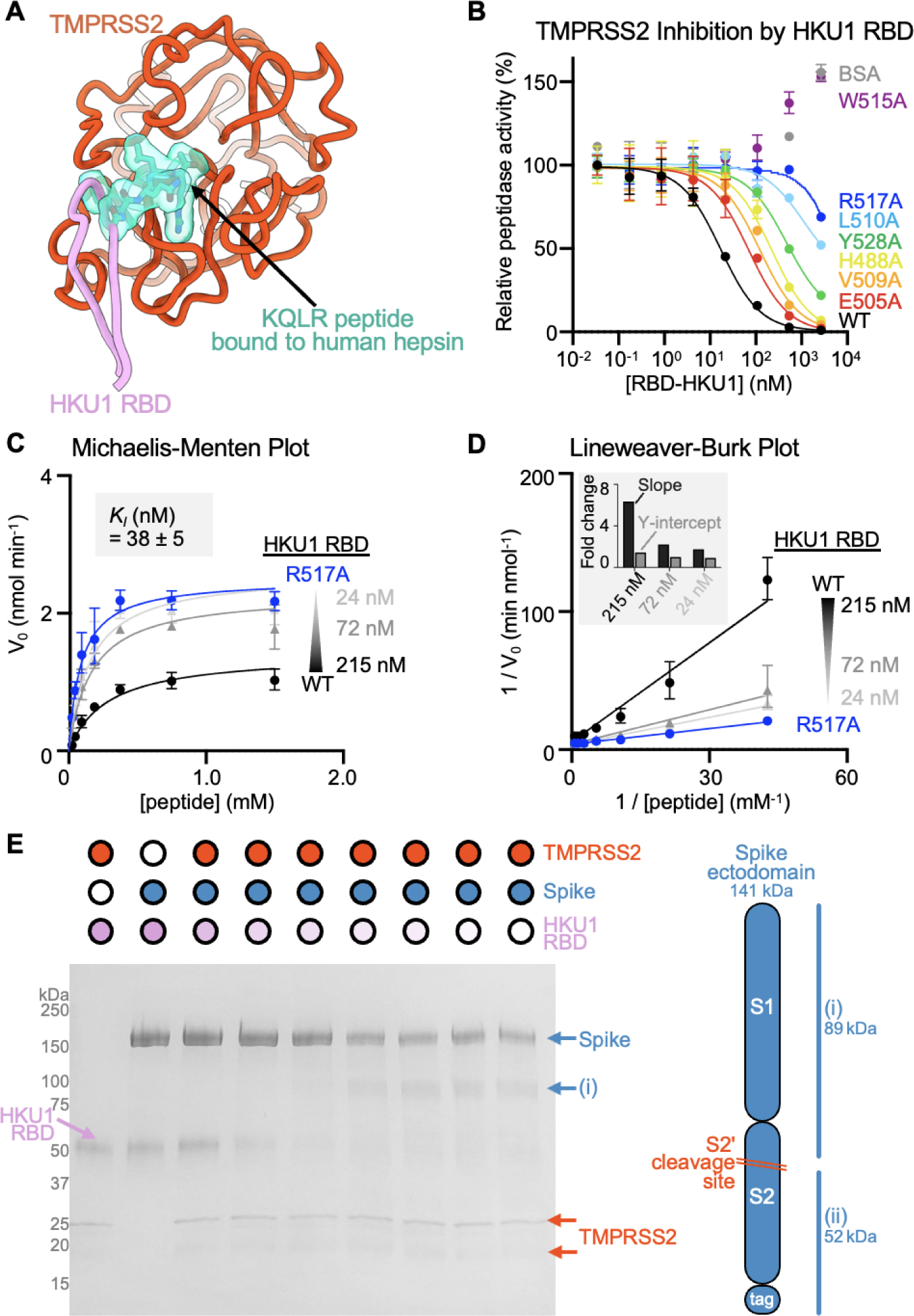
HKU1 RBD binding inhibits human TMPRSS2 activity. (A) Superimposition of the KQLR chloromethylketone-bound hepsin structure (PDB ID 1Z8G^46^) to the HKU1-bound TMPRSS2 structure showing that the ligand would be precluded sterically from binding in the active site upon attachment of the HKU1 RBD (assuming an identical binding mode of the ligand). Only the TMPRSS2 peptidase domain is shown and hepsin is omitted for clarity. (B) Assessment of inhibition of human TMPRSS2 activity by the wildtype (WT) isolate N1 and interface mutant HKU1 RBDs using the fluorescent Boc-QAR-AMC peptide substrate in the presence of 3.4 nM of the TMPRSS2 ectodomain harboring the DS, SUMO, and N249 glycan. Data are shown as the geometric mean and standard deviation of 2-3 technical replicates. (C-D) Michaelis–Menten plot (C) and Lineweaver-Burke plot (D) of initial reaction velocities in function of the concentration of fluorescent Boc-QAR-AMC peptide substrate, at 22 °C, in the presence of 5 nM TMPRSS2 and the wildtype (WT) isolate N1 or R517A interface mutant HKU1 RBDs. The inhibitor constant *K_I_*was determined through fitting competitive inhibition using GraphPad Prism. Data are shown as the geometric mean and standard deviation of three technical replicates. The inset in panel D shows the relative change in slope (*K_M_*/*V_max_*) and y axis intercept (1/*V_max_*) for the different concentrations of wildtype HKU1 RBD relative to the R517A RBD. (E) Reducing SDS-PAGE analysis of proteolytic processing at 37°C of 5 µg of SARS-CoV-2 S 2P (S2P) by 0.5 µM TMPRSS2 preincubated with various concentrations of the HKU1 isolate N1 RBD.

## Molecular basis of HKU1 specificity for TMPRSS2

Our structural data provide a blueprint to understand the HKU1 specificity for the TMPRSS2 receptor. Amino acid sequence alignment of TMPRSS2 orthologs reveals that the HKU1 binding site is mostly conserved among mammals, partially conserved in reptiles and birds, and poorly conserved in amphibians and other vertebrates (**Figure 4A-C**). We scored the compatibility with the HKU1 RBD of residues mutated relative to human TMPRSS2 using proteinMPNN^47^ and observed high probabilities for mammalian TMPRSS2 residues whereas other TMPRSS2s had lower scores. Among weakly conserved residues, mutations at positions Y414, D417, and Y469 were predicted to have the greatest reductions of HKU1 RBD binding to mammalian TMPRSS2 orthologs (**Figure 4A-C**). Moreover, mutations at positions T341, Y416, L419, and S463 were anticipated to particularly impact HKU1 RBD binding to non-mammalian TMPRSS2s.

**Figure 4.**
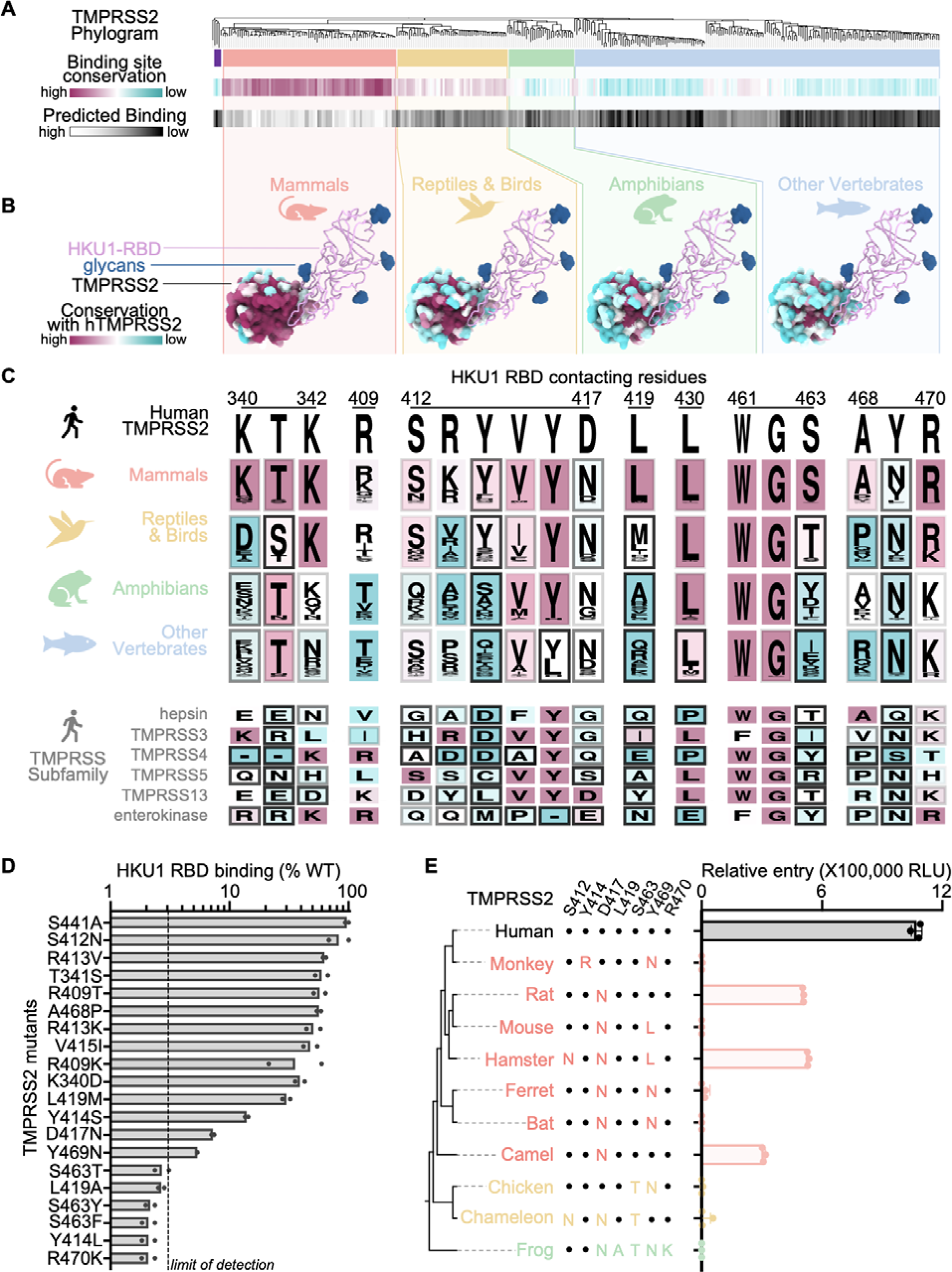
Molecular determinants of HKU1 utilization of human TMPRSS2 and host receptor tropism. (A, top) Neighbor-Joining phylogram of TMPRSS2 orthologs, clustered in red (mammals), yellow (reptiles/birds), green (amphibians), and blue (other vertebrates). The tree is rooted (far left, purple) on TMPRSS3 4, 5, and 13. (A, middle) The average amino acid sequence conservation (BLOSUM62) of the HKU1 RBD-contacting residues in each TMPRSS2 ortholog, relative to human TMPRSS2, is rendered from maroon (conserved) to cyan (not conserved). (bottom) As a proxy for predicted binding compatibility, the ProteinMPNN^47^ probability score of the HKU1 RBD-contacting residues in each TMPRSS2 ortholog was calculated and rendered from white (highest score, eg. human TMPRSS2) to black (lowest score). (B) Average amino acid sequence conservation (BLOSUM62) with human TMPRSS2 (hTMPRSS2) of residues from the TMPRSS2 clusters in (A) plotted on the surface of the human TMPRSS2 structure and rendered from maroon (conserved) to cyan (not conserved). The HKU1 RBD is shown as pink ribbons with N-linked glycans rendered as blue surfaces). (C) Logo plots of amino acids equivalent to the TMPRSS2 residues interacting with the HKU1 RBD from the ortholog clusters in (A) and from members of the TMPRSS protease subfamily colored by conservation with human TMPRSS2 from maroon (conserved) to cyan (not conserved). The shade of the frame for each residue represents the predicted binding score from proteinMPNN. (D) Binding of TMPRSS2 point mutants (corresponding to polymorphisms identified in TMPRSS2 orthologs or in humans) to the HKU1 RBD immobilized on biolayer interferometry SA biosensors. Data are shown as the area under the binding response curve normalized to the signal of WT TMPRSS2. Data are shown as the mean of two technical replicates presented in **Figure S5**. Dashed line shows the practical limit of detection. (E) HKU1 S VSV entry into HEK293T cells transiently transfected with one of the indicated TMPRSS2 orthologs. Key TMPRSS2 amino acid residues interacting with the HKU1 RBD are indicated to highlight conserved and divergent positions relative to human TMPRSS2. Data are shown as the geometric mean and standard deviation of three technical replicates.

To test this hypothesis, we expressed human TMPRSS2 harboring the K340D, T341S, R409T, S412N, R413K, Y414S, V415I, D417N, L419A/M, S463T/Y/F, A468P, Y469N, or R470K mutations and analyzed binding to the HKU1 isolate N1 wildtype RBD (**Figure 4D and Figure S5**). All mutants bound more weakly to the HKU1 RBD than wildtype TMPRSS2 with the Y414S/L, D417N, L419A, S463T/Y/F, and Y469N causing marked binding reductions, consistent with the compatibility probably score (**Figure 4C**). Contrary to its predicted score, R470K in TMPRSS2 was also severely impaired in its ability to bind the HKU1 RBD, possibly due to a requirement of R470 pi-stacking interactions with HKU1 R517 to overcome charge repulsion effects^48^. These findings likely narrow the TMPRSS2 species tropism of HKU1 as mammalian TMPRSS2s frequently harbor D417N and Y469N substitutions sometimes accompanied by mutations of residue Y414. Furthermore, S463 and L419 are not conserved for non-mammalian TMPRSS2s and R470 is only partially conserved among reptile/bird TMPRSS2s and not conserved for TMPRSS2s found in amphibians and other vertebrates. The combination of these residue substitutions along with the Y469 and Y414 mutations, are expected to limit the ability of the HKU1 RBD to engage non-mammalian TMPRSS2 orthologs.

Using a vesicular stomatitis virus (VSV) pseudotyped with HKU1 isolate N1 S, we found that transient transfection of rat, hamster and camel TMPRSS2s promote HKU1 S-mediated entry into HEK293T cells, in addition to human TMPRSS2 (**Figure 4F**). Each of these three non-human TMPRSS2 orthologs harbors the D417N residue substitution, suggesting that this mutation alone does not prevent entry. Although rat TMPRSS2 (possessing D417N) and hamster TMPRSS2 (possessing D417N, Y469L, and S412N mutations) rendered cells susceptible to HKU1 S VSV, mouse TMPRSS2 (possessing D417N and Y469L mutations) did not (**Figure 4F**). We predict that a mutation (potentially S412N) in hamster TMPRSS2 epistatically offsets a possible binding defect from Y469L. The lack of HKU1 S VSV entry into cells transiently transfected with ferret, greater horseshoe bat, and African green monkey TMPRSS2s is rationalized by the reduced HKU1 RBD binding of the TMPRSS2 Y469N and D417N mutants and likely the Y414R mutant. As anticipated for TMPRSS2 orthologs harboring S463T, R470K, or L419A mutations, non-mammalian (chicken, African clawed frog, and chameleon) TMPRSS2s did not support HKU1 S VSV entry. Collectively, our data explain the efficient HKU1 use of human TMPRSS2 as entry receptor and reveal that rat, hamster and camel TMPRSS2 orthologs support HKU1 S-mediated entry into cells.

## Functional consequences of TMPRSS2 site mutants in humans

We found several low frequency TMPRSS2 single nucleotide polymorphisms (SNPs) at residue positions involved in HKU1 binding and several of them correspond to missense mutations observed in TMPRSS2 paralogs and orthologs (**Figure 4C and Data S1-S2**). Among the SNPs we characterized experimentally, some of the mutations reduced binding to the HKU1 RBD (T341S, R409T/K, S412N, V415I and A468P) whereas others almost entirely abrogated binding (S463F and R470K). The most frequently sampled SNP is V415I (rs148125094, allele frequency: 1.1x10^-3^) which led to reduced but detectable binding to the HKU1 isolate N1 RBD relative to wildtype human TMPRSS2 (**Figure 4D, Figure S5 and Data S1-S2**). The S463F SNP (rs2146419758, allele frequency: 2.1x10^-6^), however, abrogated binding to the wildtype HKU1 RBD, concurring with the fact that none of the energetically favored phenylalanine side chain rotamers at this position would be sterically compatible with the binding interface in our cryoEM structure (**Figure 4D, Figure S5 and Data S1-S2**). In summary, although several TMPRSS2 SNPs map to the site of HKU1 attachment, some of them (e.g. S463F and R470K) possibly reducing susceptibility to HKU1 infection, the low frequency of TMPRSS2 SNPs is unlikely to majorly impact HKU1 transmission in the human population.

Residues involved in binding to the HKU1 RBD, markedly diverge across human type 2 transmembrane serine proteases with G462 being the sole residue strictly conserved among them (**Figure 4C**). Members of the TMPRSS protease subfamily harbor non-conservative mutations at several positions involved in HKU1 binding relative to TMPRSS2. This includes L419A (TMPRSS5), S463T/Y (hepsin, TMPRSS4, TMPRSS13, and enterokinase), and Y469N (TMPRSS3, TMPRSS5, TMPRSS13, and enterokinase), which we found to abrogate TMPRSS2 binding to the HKU1 RBD (**Figure 4C-D and Figure S5**). These data therefore rationalize the lack of HKU1 S-mediated fusion observed in the presence of TMPRSS3, TMPRSS4, and TMPRSS13^29^ and explain the HKU1 specificity for TMPRSS2.

## Glycan shielding and conformational masking mediates immune evasion

Although the HKU1 RBM protrudes out from the closed S trimer, leading to markedly increased solvent exposure relative to sarbecovirus RBMs^12, 49^, TMPRSS2 binding would be precluded due to steric hindrance involving both proteinaceous and oligosaccharide moieties from neighboring protomers (**Figure 5A-B**). Sialoglycan receptor binding, which is known to promote RBD opening^30^, would relieve these steric constraints and enable TMPRSS2 attachment to an RBD in the open conformation (**Figure 5C**). Similar to other coronaviruses, this cascade of events is expected to lead to fusion of the viral and host membranes to promote genome delivery to a target cell and initiation of infection^49–51^.

**Figure 5.**
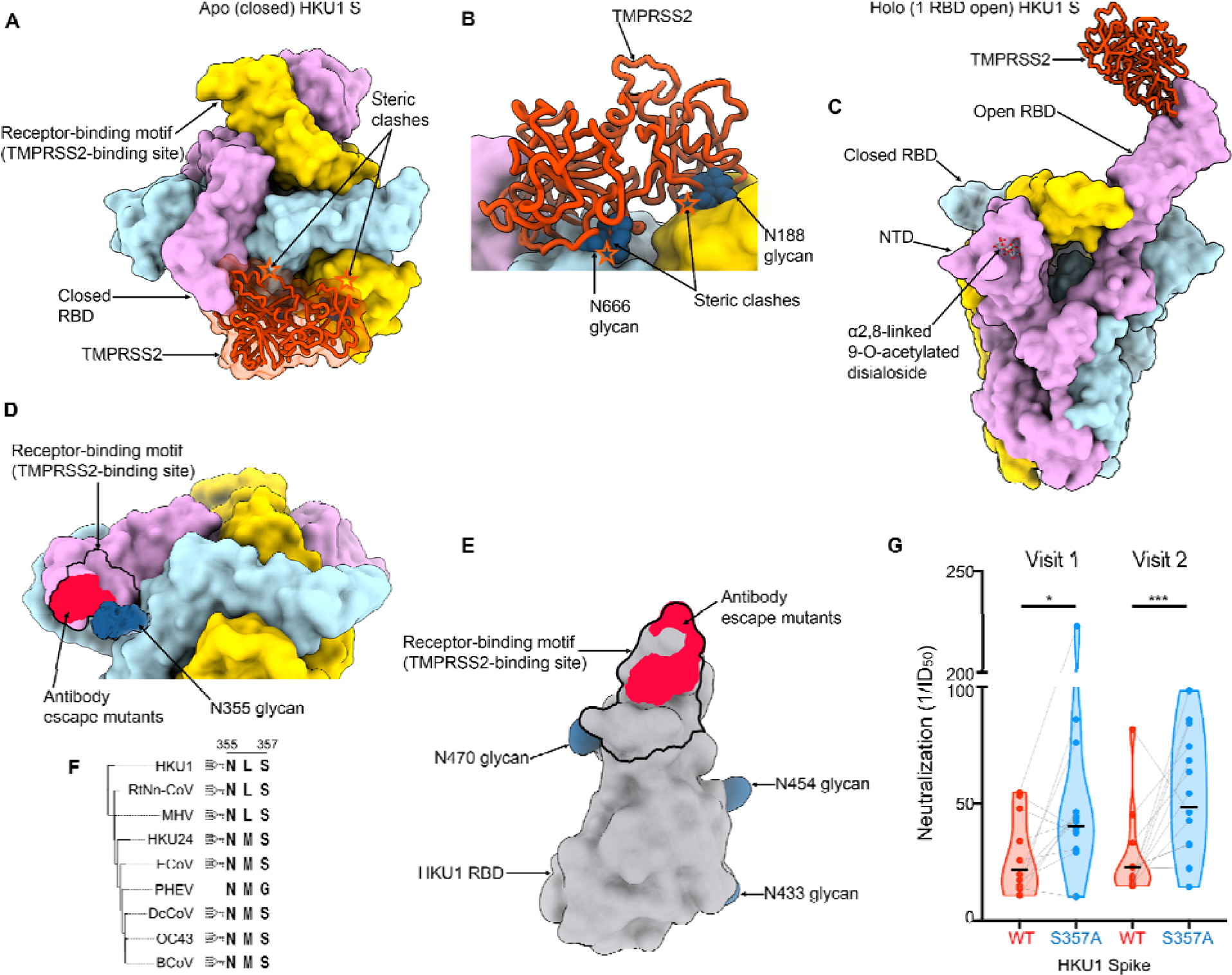
Glycan shielding and conformational masking mediate HKU1 immune evasion. (A-B) Composite model obtained by superimposing the TMPRSS2 (orange)-bound HKU1 RBD cryoEM structure onto the apo closed HKU1 S trimer structure (PDB 8OHN)^30^ rendered as a surface with each protomer colored distinctly viewed in two orthogonal orientations. In the closed trimer conformation, TMPRSS2 binding would be hindered by steric clashes (orange stars) with neighboring protomers, including with N-linked glycans N188, N355, N666 (shown as blue spheres). (C) Composite model obtained by superimposing the TMPRSS2 (orange)-bound HKU1 RBD cryoEM structure onto the α2,8-linked 9-O-acetylated disialoside-bound HKU1 S trimer with one open RBD (PDB 8OPN)^30^ engaging the TMPRSS2 receptor. (D) Zoomed-in view of the apo closed HKU1 S trimer structure (PDB 8OHN)^30^ showing the unsharpened cryoEM density for the N355 glycan partially masking the receptor-binding motif (black outline) and the site of vulnerability to the mHKUS-2 and -3 neutralizing antibodies^45^ (neon red). The extent of the N355 glycan-mediated glycan shielding is expected to be even greater than shown as only the first three monosaccharides are resolved in the map. (E) HKU1 RBD rendered as a grey surface with the TMPRSS2 footprint delineated with a black outline and the escape mutants (D511,H512,W515,R517) identified for the neutralizing antibodies mHKUS-2 and -3 shown in neon red. The HKU1 RBD N-linked glycans are rendered as blue surfaces. (F) Neighbor-Joining phylogram of embecovirus S glycoproteins focused on the HKU1 N355 glycosylation site. Included S sequences are HKU1 (AZS52618.1), RtNn-CoV (RtNn-CoV/SAX2015, ATP66762.1), mouse hepatitis virus (MHV, C0KYY9), HKU24 (A0A866W1F1), Equine coronavirus (ECoV, E5RPZ2), Porcine hemagglutinating encephalomyelitis virus (PHEV, A0A1Z2WUW0), Dromedary Camel coronavirus (DcCoV, HKU23, A0A291L0R6), OC43 (P36334) and bovine coronavirus (BCoV, P15777). (G) Neutralization of VSV pseudotyped with wildtype isolate N1 HKU1 S (WT) or the N355 glycan knockout (S357A) mutant by serum antibodies elicited by HKU1 infection. Visits 1 and 2 correspond to blood draws at the time of PCR-positive testing and 1-2 months later, respectively^52, 53^. Neutralization values were compared by two-way ANOVA with the Geisser-Greenhouse correction and Tukey’s multiple comparison test; * corresponds to a P value of 0.02, while *** corresponds to a P value of 0.0007.

Strikingly, the HKU1 glycan at position N355 from a neighboring RBD masks the RBM (**Figure 5D**), as observed in an apo closed HKU1 S structure^30^, sterically limiting RBM accessibility in this conformation. As a result, residues D511, H512, W515 and R517, which map to the RBM and contribute to the epitopes recognized by the previously described mHKUS-2 and -3 neutralizing monoclonal antibodies^45^, are also partially masked by the HKU1 N355 glycan which would reduce targeting by this class of antibodies (**Figure 5D-E**). Given the overlap between the epitopes of the mHKUS-2 and -3 antibodies and the RBM, the primary mechanism of neutralization for these antibodies is most likely through direct competition with HKU1 attachment to the TMPRSS2 host receptor. Given the conservation of this glycan in most embecovirus S glycoproteins (**Figure 5F**), similar masking is anticipated to occur, as confirmed in the prefusion OC43 S structure^26^ although the role of the topological equivalent of the RBD (domain B) remains unclear for many members of this subgenus.

To investigate the relationship between antibody specificity and viral inhibition, we assessed neutralizing antibody titers against an HKU1 isolate N1 S VSV using human sera collected after PCR-confirmed HKU1 infection. These serum samples were collected in four European countries between 2007 and 2010 at the time of PCR-positive testing (Visit 1) and 1-2 months later (Visit 2)^52, 53^. Neutralizing antibody titers ranged from half-maximum inhibition dilution of 1/10 to 1/80 for a geometric mean of 1/23 for visit 1 and 1/49 for visit 2 (**Figure 5G and Figure S6**). Serum neutralizing activity was enhanced upon removal of the N355 glycan (via S357A mutation), relative to the wildtype HKU1 S, concurring with the partial masking of the RBM by this oligosaccharide (**Figure 5G and Figure S6**). These findings indicate that RBM-directed serum antibodies are a key component of neutralization, in line with prior studies of potent sarbecovirus and merbecovirus antibodies^49, 54–59^. Collectively, our data reveal that HKU1 relies on conformational masking and glycan shielding to balance conflicting requirements for immune evasion and receptor engagement, ultimately modulating viral fitness.

## Discussion

HKU1 genomes sequences are divided into three genotypes designated A, B and C with evidence of recombination among them^5, 9^. The S glycoproteins of genotypes A and B share 85% amino acid sequence identity whereas that of genotypes B and C are almost identical^29^. Furthermore, HKU1 RBDs are highly divergent with ∼74% amino acid sequence identity between the HKU1 isolate N1 RBD (genotype A) and the isolate N2 RBD (genotype B) or the isolate N5 RBD (genotype C). This magnitude of sequence divergence is comparable to that observed among the RBDs of distinct sarbecovirus clades^60, 61^ and may participate in explaining that protective immunity to seasonal coronaviruses is of short duration due to antigenic drift^62, 63^. Furthermore, HKU1 infections elicit modest neutralizing antibody titers (at least in the cohort analyzed here), as compared to SARS-CoV-2 infection^64^, which may further compound the durability and breadth of humoral immune responses. The enhanced neutralization of an HKU1 S mutant lacking the N355 glycan, relative to wildtype, with a panel of human sera collected after HKU1 infection show that RBM-directed antibodies are making a key contribution to neutralizing activity. These findings may explain the observed polymorphism of TMPRSS2-interacting residues among HKU1 isolates and are reminiscent of observations made for SARS-CoV-2^55, 65, 66^ and 229E^63, 67^. Our structural data therefore provide a molecular blueprint to follow evolution of RBM residues and their impact on receptor binding and immune evasion. The identification of the RBM as a key site of vulnerability further suggests that removal of the N355 glycan could be a suitable vaccine design strategy to improve the potency of antibody responses elicited by HKU1 S and for most viruses from the embecovirus subgenus. Although the OC43 domain B is not known to be a functional RBD, potent neutralizing antibodies recognizing this region were described^68^, underscoring the possible key role of this conserved embecovirus glycan for immune evasion. Since HKU1 recognizes 9-O-acetylated sialosides using the S NTD^27, 30, 69, 70^, similar to OC43 and bovine coronavirus^19, 26^, the isolation of a neutralizing antibody competitively inhibiting sialoside binding to the OC43 NTD^68^ suggests that this mechanism of action is likely conserved against many, if not all, lineage A β-coronaviruses attaching to cell-surface carbohydrates. Future studies will elucidate the importance and possible synergy of these different types of antibodies, along with that of non-neutralizing antibodies, for in vitro neutralization and in vivo protection against these viruses.

Rapid antigenic evolution, masking of epitopes and exposure of immune-dominant epitopes targeted by non-neutralizing antibodies are immune-evasion strategies widely used by viruses. Coronavirus S trimers can adopt at least two distinct conformations of their RBDs enabling to mask or expose the RBM in the closed and open states, respectively. Although the SARS-CoV-2, SARS-CoV-1 and MERS-CoV S trimers appear to populate this conformational landscape spontaneously^12, 49, 71, 72^, most coronavirus S trimers appear to preferentially adopt a closed conformation^18, 26, 51, 73–79^. Recent work showed that α2,8-linked 9-O-acetylated disialosides binding to the HKU1 S NTD induces conformational changes leading to RBD opening^30^, which our data reveal to expose the TMPRSS2-binding site. This elegant mechanism would allow to ensure proper spatial and temporal coordination of these conformational changes, ensuring that RBD opening occur upon attachment to a target cell, thereby balancing the conflicting requirements for receptor binding to initiate infection (which requires RBD opening) and immune evasion through masking of a key site of vulnerability (RBM) recognized by polyclonal serum neutralizing antibodies (which requires RBD closing).

Although HKU1 has been proposed to have originated in rodents, due to phylogenetic clustering with rodent coronaviruses^80^, evidence supporting or invalidating this hypothesis is still lacking. We identified the molecular determinants of HKU1 utilization of TMPRSS2 and revealed that rat, hamster and camel, but not mouse, TMPRSS2 orthologs support HKU1 S-mediated entry into cells. These data are compatible with a possible rodent origin of HKU1 and the putative involvement of the species identified here as reservoir or intermediate hosts. We note, however, that host tropism is a complex process involving additional factors than receptor recognition, including proteolytic spike activation and innate immune antagonism, for a successful infection to occur. Considering the wide use of golden syrian hamsters as small animal models^59, 81, 82^, our data suggest a possible path to develop an HKU1 challenge model to study pathogenicity, correlates of protection and the effectiveness of countermeasures.

## Acknowledgements

This study was supported by the National Institute of Allergy and Infectious Diseases (P01AI167966, DP1AI158186 and 75N93022C00036 to D.V.), a Pew Biomedical Scholars Award (D.V.), an Investigators in the Pathogenesis of Infectious Disease Awards from the Burroughs Wellcome Fund (D.V.), the University of Washington Arnold and Mabel Beckman cryoEM center and the National Institute of Health grant S10OD032290 (to D.V.). D.V. is an Investigator of the Howard Hughes Medical Institute and the Hans Neurath Endowed Chair in Biochemistry at the University of Washington. We gratefully acknowledge the Biobank Antwerp, Belgium (ID: BE 71030031000) for providing the human sera collected in the GRACE study (www.grace-lrti.org) supported by the 6th Framework Program of the European Commission, contract no. LSHM-CT-2005-518226).

## Author Contributions

MM, YJP, and DV conceived the study and designed the experiments; MM designed the engineered TMPRSS2 constructs and carried out activity assays. JB performed the S2P cleavage assay. MM designed and cloned protein constructs. MM, CS and JB recombinantly expressed and purified glycoproteins. CS performed binding assays. Y.J.P. carried out cryoEM specimen preparation, data collection and processing. Y.J.P. and D.V built and refined atomic models. EW and AT carried out the single nucleotide polymorphism analysis. G.I. provided unique reagents. MM, YJP, and D.V. analyzed the data and wrote the manuscript with input from all authors. DV supervised the project.

## MATERIALS AND METHODS

### Cells

Cell lines used in this study were obtained from ATCC (HEK293T and VeroE6) or Thermo Fisher Scientific (ExpiCHO-S cells and Expi293F cells).

### Constructs

The membrane-anchored HKU1 isolate N1 S (ref. seq. YP_173238, genotype A) with a 21 amino acid C-terminal deletion and six residues flexible linker followed by C-terminal HA tag were codon-optimized, synthesized, and inserted the pcDNA3.1(+) vector by Genscript. The HKU1 RBD constructs encoding S residues 320-614 of the wildtype isolate N1 (ref. seq. YP_173238, genotype A) and the H488A, E505A, V509A, L510A, W515A, R517A and Y528A HKU1 RBD interface mutants, containing N-terminal mu-phosphatase secretion signal peptide, with or without an avi tag followed by three residues flexible linker and C-terminal octa-histidine tag were codon optimized, synthesized, and inserted into the pcDNA3.1(+) vector by Genscript. The membrane-anchored TMPRSS2 and TMPRSS2 ectodomain constructs were codon-optimized, synthesized, and inserted into a pCMVR vector. TMPRSS2 ectodomain constructs were further mutated by In-Fusion Assembly (Takara Bio). All TMPRSS2 sequences are listed in **Table S1**.

The SARS-CoV-2 S 2P (Wuhan-Hu-1) ectodomain construct^12^ harbors an N-terminal mu-phosphatase secretion signal peptide, S1/S2 cleavage site R682S/R683G/R685G mutations, K986P/V987P mutations, one residue linker followed by C-terminal hexa-histidine tag and foldon trimerization domain was codon-optimized, synthesized, and inserted the pCMV vector by Genscript.

### Generation of pseudoviruses

To produce pseudoviruses for entry and neutralization assays, HEK293T cells were seeded in DMEM enriched with 10% FBS (Hyclone), 1% Penstrep at the appropriate density to yield 80% confluency in polylysine-coated 100 mm cell culture dishes and placed in an incubator at 37°C with 5% CO_2_. After 18-22 hr incubation, cells were washed with OPTI-MEM (Life Technologies). 24µg of either the wildtype HKU1 isolate N1 (genotype A) S or the S357A mutant S plasmids were prepared in 1.5mL of OPTI-MEM and combined with 60µL of Lipofectamine 2000 (Life Technologies) diluted in 1.5mL of OPTI-MEM and incubated at room temperature for 15-20 min. The mixture was added to the HEK293T cells which were placed for 2 hours in an incubator at 37°C with 5% CO_2_ after which 2mL of DMEM enriched with 20% FBS and 2% PenStrep was added to the transfected cells and incubated overnight. The following day, cells were washed with DMEM and transduced with VSVΔG/Fluc and incubated for 2 hours at 37°C with 5% CO_2_. After washing with DMEM, medium supplemented with anti-VSV-G antibody (I1-mouse hybridoma supernatant diluted to 1:25 from CRL-2700, ATCC) was added to the cells to reduce background from the parental virus and an additional incubation at 37°C with 5% CO_2_ was performed overnight. The next day, the supernatant from the cells was harvested from the 100 mm dishes, further clarified by centrifugation at 3,000xg for 10 minutes, filtered (0.45μm), and concentrated 10 times by using centrifugal devices with 30 kDa cutoff membranes. Pseudoviruses were then aliquoted and frozen at -80°C until used.

### Pseudovirus neutralization and entry assays

HEK293T cells were cultured in DMEM with 10% FBS (Hyclone), 1% PenStrep and placed in cell-culture grade, polylysine-coated 100 mm plates overnight in an incubator at 37°C with 5% CO_2_. The following day, when cells reached 90% confluency, they were transfected with 4 µg of TMPRSS2 and incubated for 5 hours. The cells were subsequently trypsinized, counted and reseeded at ∼40,000 cells per well in poly-lysine-coated 96 well plates and placed in an incubator at 37°C with 5% CO_2_ overnight. The next day, a half-area 96-well plate was prepared with a 1:3 serial dilution of sera in DMEM in 22 μL final volume. 22 μL of pseudovirus was then added to each well and incubated at room temperature for 30-45 min. The media was removed from transfected HEK293T cells and 40 μL of the sera/pseudovirus mixture was added to the cells and incubated for 2 h at 37°C with 5% CO_2_ before adding 40 μL of 20% FBS and 2% PenStrep containing DMEM. No sera were used for assessment of entry with the panel of TMPRSS2 orthologs. Following 18-22 hour incubation, 40 μL of One-GloEX (Promega) was added to the cells and incubated in the dark for 5 min prior to reading on an Agilent BioTek Neo2 plate reader. Relative luciferase units were plotted and normalized in Prism (GraphPad) using a zero value of cells alone and a 100% value of 1:2 virus alone. Nonlinear regression of log(inhibitor) vs. normalized response was used to determine ID_50_ values from curve fits. At least two biological replicates with two distinct batches of pseudovirus were conducted for each serum sample.

### Recombinant glycoprotein production

The SARS-CoV-2 S 2P ectodomain, HKU1 RBDs or TMPRSS2 ectodomains were produced and purified using Expi293F or ExpiCHO-S cells. Expi293F cells were grown to a density of 3 x 10^6^ cells/mL and transfected using the ExpiFectamine 293 Transfection Kit (ThermoFisher Scientific) and expression carried out for 3 to 5 days post-transfection at 37°C with 8% CO_2_. ExpiCHO-S cells were grown to a density of 6 x 10^6^ cells/mL and transfected using the ExpiFectamine CHO Transfection Kit (ThermoFisher Scientific) and expression carried out for 7-10 days post-transfection at 37°C with 8% CO_2_. The SARS-CoV-2 S 2P ectodomain and HKU1 RBDs were purified from clarified supernatants using HisTrap HP affinity columns (Cytiva) and washed with ten column volumes of 10 mM imidazole, 25 mM sodium phosphate pH 8.0, and 300 mM NaCl before elution with two column volumes of 300 mM imidazole, 25 mM sodium phosphate pH 8.0, and 300 mM NaCl. The purified HKU1 RBD proteins were buffer exchanged into 20 mM sodium phosphate pH 8 and 100 mM NaCl or 50 mM Tris pH 8.0 and 150 mM NaCl. The HKU1 RBDs were biotinylated using the BirA biotin ligase reaction kit (Avidity) following the manufacturer’s protocol. Biotinylation was carried out at room temperature for 30 minutes followed by incubation for 10 hours at 4°C. TMPRSS2 constructs were purified using TALON metal affinity resin (Takara Bio) and washed with 100 column volumes of 5 mM imidazole, 50mM HEPES pH 7.5, and 150mM NaCl prior to elution with 600 mM imidazole, 50mM HEPES pH 7.5, and 150mM NaCl. Purified TMPRSS2s were diluted to 0.3 mg/ml and then digested with a 1/500 dilution of enterokinase (EKmax, ThermoFisher) for 16 hours at room temperature or 1 hour at 37 °C to convert the zymogen in active enzyme, remove the SUMO tag and the His tag. Digested TMPRSS2 was then passed through fresh TALON metal affinity resin to remove enterokinase and the cleaved His tag. Active TMPRSS2 with the DS mutation, SUMO tag, and restored N249 glycan activated itself during expression and purification, and did not require enterokinase activation. Purified HKU1 RBD and TMPRSS2 – except for TMPRSS2 used for Figure 4D – were then further purified by size exclusion chromatography using a Superdex 200 Increase 10/300 GL column (Cytiva) and concentrated using centrifugal filters (Amicon Ultra) before being flash frozen.

### Biolayer interferometry

Biotinylated HKU1 wildtype (isolate N1, genotype A) RBD and H488A, E505A V509A, L510A, W515A, R517A and Y528A HKU1 RBD interface alanine mutants were diluted to concentrations of 0.002 mg/mL in modified kinetics buffer (1% BSA, 0.06% Tween20, 1x Cold Spring Harbor PBS pH 7.4) and loaded onto pre-hydrated streptavidin biosensors to a 1 nm total shift. The loaded tips were dipped into 100 nM TMPRSS2 (harboring the S441A residue substitution and the C379-T447C disulfide bond) for 200 or 300 seconds followed by dissociation in modified kinetics buffer for 500 seconds. Analysis of TMPRSS2 interface alanine mutants was carried out similarly but with constructs harboring S441 and the C379-T447C disulfide bond. Binding curves were plotted using Graphpad Prism 10.1.2. The area under the curves were determined by summation of the binding response values for each reading, with a limit of binding detection of 0.05 nm. Sums were normalized to WT, as was the limit of detection which corresponded to 2 ± 0.5 % (the practical limit of binding detection was set to two standard deviations above this value, i.e. 3.1 % of WT binding).

### TMPRSS2 enzymatic activity assay

Peptidase assays were performed with Boc-QAR-AMC substrate (GlpBio) in black half area 96 well plates (Greiner Bio-One Fluotrac) in 100 µl reaction volumes in a Agilent BioTek Microplate Reader at 22 °C, monitoring fluorescence every two minutes over 30 minutes at 341 nm excitation and 441 nm emission. The slope of the fluorescence curve over the first 10 minutes (corresponding to <5 % substrate conversion) was used for velocity calculations. AMC concentration was calculated using standard curves at each substrate and HKU1 RBD concentration, to correct for the inner filter effect. For determination of TMPRSS2 enzyme kinetics, serial dilutions of Boc-Gln-Ala-Arg-AMC substrate were reacted with 6.8 nM TMPRSS2. For determination of the *K_I_*of the HKU1 RBD, 5 nM TMPRSS2 was reacted with serial dilutions of Boc-QAR-AMC substrate in the presence of 215, 72, or 24 nM HKU1-RBD (TMPRSS2 was incubated for 2 minutes with the HKU1 RBD at 22 °C prior to the addition of substrate). Velocities were plotted and fit using GraphPad Prism (v10.1.1).

### Spike Cleavage Assay

Spike cleavage assays were performed with SARS-CoV-2 S-2P ectodomain in 20mM phosphate pH 8.0 and 100mM NaCl incubated with 500nM TMPRSS2 in 50 mM HEPES pH 7.5 150mM NaCl. Samples were diluted in 50 mM HEPES pH 7.5 150mM NaCl, mixed, and incubated at 37°C with 300 rpm shaking. Reactions were stopped at various time points by mixing with 4x gel loading buffer containing 200mM Tris pH 6.8, 8% SDS, 20% β-mercaptoethanol, 40% Glycerol, 0.2% Bromophenol blue. The HKU1 RBD inhibition assays were performed by incubating the HKU1 RBD (without avi tag) with TMPRSS2 (harboring the N-terminal SUMO fusion and the C379-T447C disulfide bond) in 50 mM HEPES pH 7.5 150mM NaCl for 5 minutes at 37°C prior to adding SARS-CoV-2 S 2P and incubating at 37°C with shaking at 600 rpm for 15 minutes prior to stopping the reaction by mixing with 4x gel loading buffer containing 200mM Tris pH 6.8, 8% SDS, 20% β-mercaptoethanol, 40% Glycerol, 0.2% Bromophenol blue. Samples were then analyzed by SDS-PAGE.

### Human serum sample donors

The patients/participants provided their written informed consent to participate in this study which was reviewed and approved by Antwerp University Hospital. Human sera were collected after PCR-confirmed HKU1 infection in four European countries between 2007 and 2010 at the time of PCR-positive testing (Visit 1) and 1-2 months later (Visit 2). Demographic data can be found in **Data S3**.

### Single nucleotide polymorphism (SNP) analysis

The SNP information was extracted from two human population allele frequency database, <u>GnomAD (version 4)</u> and <u>Regeneron’s Million Exome Variant database</u>. For GnomAD v4, we utilized the allele frequencies of short genetic variations from the exome sequencing data of 730,947 individuals. These genetic variations have passed the QC (i.e. “FILTER” column in the VCF = “PASS”). GnomAD v4 provided the functional consequences of the genetic variations from the <u>VEP tool</u>. We obtained the predicted deleteriousness of mutations from AlphaMissense^83^. For Regeneron’s Million Exome Variant data, we utilized the allele frequencies of short genetic variations from the exome sequencing data of 983,578 individuals. We observed a total of 49 short genetic variations in the GnomAD v4 and the Regeneron’s Million Exome Variant databases.

### CryoEM sample preparation, data collection and data processing

Complex formation was performed by mixing a 1:1.2 molar ratio of the human TMPRSS2 ectodomain (harboring the S441A residue substitution and a C379-T447C disulfide bond) with the HKU1 isolate N1 RBD (residues 320-614 with C-terminal octa-histidine tag and no avi tag) before incubation for 1 hour at 4°C. CryoEM grids of the complex were prepared using three separate methods and data were combined during data processing. For the first dataset, 3 µL of 1.1 mg/mL or 0.5 mg/mL of complex were loaded onto freshly glow discharged R 2/2 UltrAuFoil grids^84^ prior to plunge freezing using a Vitrobot Mark IV (ThermoFisher Scientific) with a blot force of 0 and 6 sec blot time at 100% humidity and 22°C. 6,681 movies from UltrAuFoil grids with 1.1 mg/mL protein complex were collected with a defocus range comprised between - 0.2 and -3 μm and stage tilt angle of 0° and 20°^85^. 3,820 movies from UltrAuFoil grids with 0.5 mg/mL protein complex were collected with a defocus range comprised between -0.2 and -3 μm and stage tilt angle of 30° and 45°. For the second dataset, 3 µL of 1 mg/mL complex was added to the glow discharged side of R 2/2 UltrAuFoil grids and 1µL was added to the back side before plunging into liquid ethane using a GP2 (Leica) with 6 sec blot time. 9,883 movies were collected with a defocus range comprised between -0.2 and -3 μm and stage tilt angle of 0° and 30°. For the third dataset, 3 µL of 8 mg/ml complex with 3 mM CHAPSO or 0.01% fluorinated octyl-maltoside (FOM) (Anatrace) were applied onto freshly glow discharged R 2/2 UltrAuFoil grids prior to plunge freezing using a vitrobot MarkIV (ThermoFisher Scientific) with a blot force of 0 and 5 sec blot time at 100% humidity and 22°C. 6,822 and 1,859 movies were collected from UltrAuFoil grids with CHAPSO and FOM detergents, respectively, with a defocus range comprised between -0.2 and -3.5 μm. The data were acquired using an FEI Titan Krios transmission electron microscope operated at 300kV and equipped with a Gatan K3 direct detector and Gatan Quantum GIF energy filter, operated in zero-loss mode with a slit width of 20 eV. Automated data collection was carried out using Leginon^86^ at a nominal magnification of 105,000× with a pixel size of 0.835 Å. The dose rate was adjusted to 15 counts/pixel/s, and each movie was acquired in super-resolution mode fractionated in 75 frames of 40 ms. Movie frame alignment, estimation of the microscope contrast-transfer function parameters, particle picking, and extraction were carried out using Warp^87^. Particles were extracted with a box size of 168 pixels with a pixel size of 1.67Å. Two rounds of reference-free 2D classification were performed using CryoSPARC^88^ to select well-defined particle images from each dataset, which were subsequently combined. Initial model generation was done using ab-initio reconstruction in cryoSPARC and used as references for a heterogenous 3D refinement in cryoSPARC. Particles belonging to classes with the best resolved HKU1 RBD and TMPRSS2 density were selected. To improve particle picking further, we trained the Topaz^89^ picker on Warp-picked particle sets belonging to the selected classes after heterogeneous 3D refinement. The particles picked using Topaz were extracted and subjected to 2D classification using cryoSPARC, which improved the number of unique 2D views. The two different particle sets picked from Warp and Topaz were merged and duplicate particle picks were removed using a minimum distance cutoff of 60Å. After two rounds of heterogeneous refinements and removal of junk particles, 3D refinement was carried out using non-uniform refinement with per-particle defocus refinement in cryoSPARC^90^ and the particles were transferred from cryoSPARC to Relion using pyem (https://github.com/asarnow/pyem) to be subjected to the Bayesian polishing procedure implemented in Relion^91^ during which particles were re-extracted with a box size of 280 pixels and a pixel size of 1.0 Å. Subsequent 3D refinement used non-uniform refinement along with per-particle defocus refinement in cryoSPARC to yield the final reconstruction at 2.9 A□ resolution comprising 810,357 particles. Reported resolutions are based on the 0.143 gold-standard Fourier shell correlation (FSC) criterion^92, 93^. Local resolution estimation, filtering, and sharpening were carried out using cryoSPARC.

### Model building and refinement

UCSF Chimera^94^ was used to rigid-body dock models into the sharpened cryoEM map and adjustments and refinement were carried out with Coot^95^ and Rosetta^96, 97^ using sharpened and unsharpened maps. Validation used Molprobity^98^, Phenix^99^ and Privateer^100^.

### Data Availability

The cryoEM map and atomic model have been deposited to the EMDB and PDB with accession codes EMD-43224 and PDB 8VGT. All datasets generated and information presented in the study are available from the corresponding authors on reasonable request. Materials generated in this study can be available on request and may require a material transfer agreement.

---

**Figure S1.**
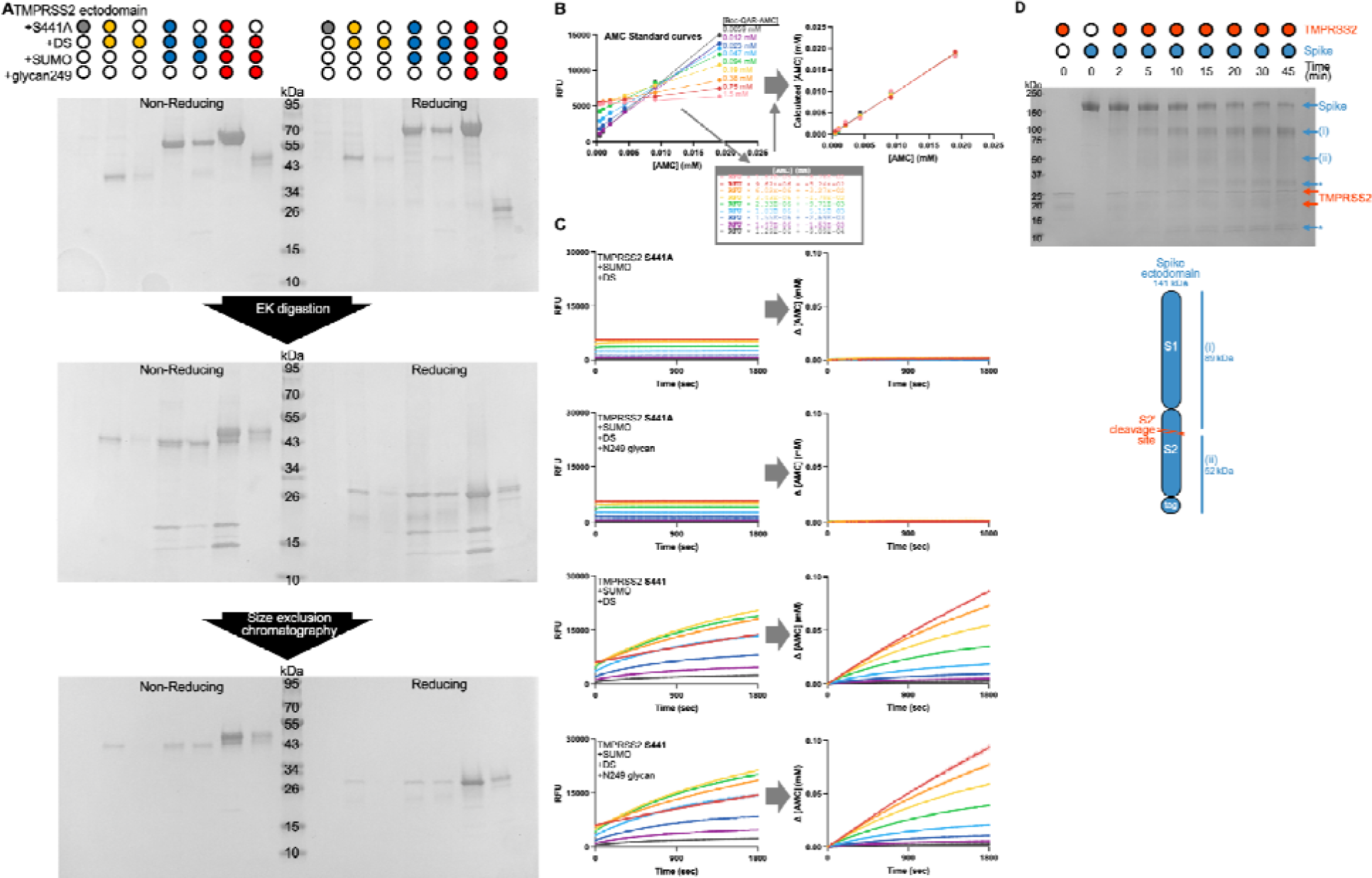
Design and functional characterization of TMPRSS2 ectodomain constructs. (A) Reducing and non-reducing SDS-PAGE analysis of the samples shown in Figure 1B, including the intermediate step between enterokinase (EK) digestion and size-exclusion chromatography (SEC) purification. (B) Summary of AMC standard curves used to calculate AMC release from the Boc-QAR-AMC peptide substrate accounting for the inner filter effect. The left panel shows the relationship between AMC concentration and measured fluorescence (RFU). The lower inset shows the numerical equations of the linear relationships shown in the top panel. The right panel shows the calculated AMC concentration for each known AMC concentration shown in the left panel, using the equations in the lower inset at the indicated Boc-QAR-AMC concentrations. (C) Raw data and calculations used for Figure 1C; the color key is identical to Figure S1B. The left panels show the raw RFU over time for TMPRSS2 with and without the S441A mutation or the restored N249 glycan, while the right panel shows the change in calculated AMC concentration over time, using the equations from Figure S1B. (D) Reducing SDS-PAGE used for analysis shown in Figure 1D. Expected products (i) and (ii), based on TMPRSS2 cleavage at the S_2_’ site, are labeled, along with additional cleavage products (asterisks). Reaction progress was monitored by densitometry of the S peak only.

**Figure S2.**
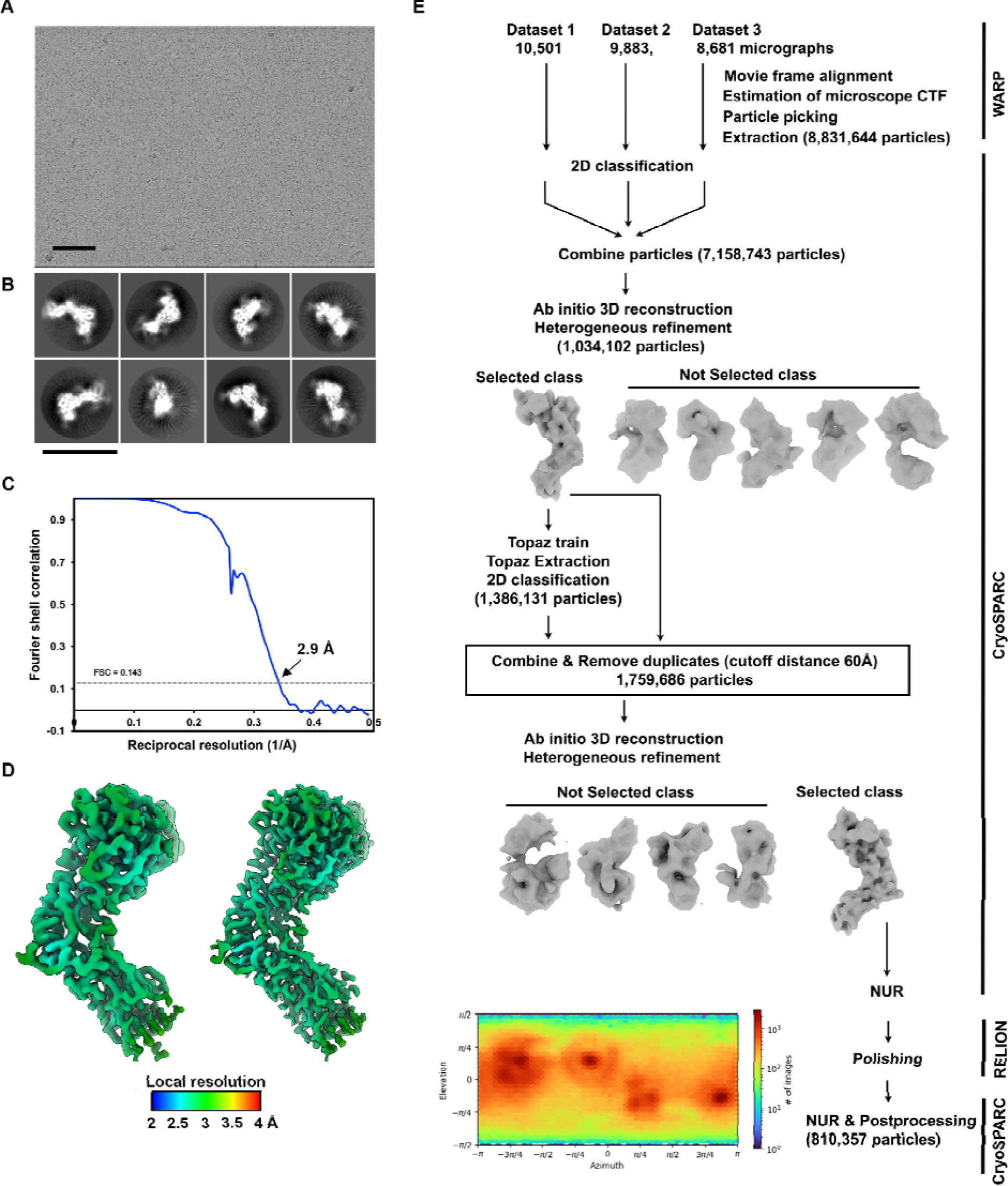
CryoEM data processing of the TMPRSS2-bound HKU1 RBD dataset. (A-B) Representative electron micrograph and 2D class averages of the TMPRSS2-bound HKU1 RBD complex embedded in vitreous ice. The scale bars represent 100 nm and 150Å respectively. (C) Gold-standard Fourier shell correlation curve. The 0.143 cutoff is indicated by a horizontal dashed line. (D) Local resolution estimation calculated using cryoSPARC and plotted on the unsharpened (left) and sharpened (right) maps. (E) Data processing flowchart. CTF: contrast transfer function; NUR: non-uniform refinement. The angular distribution calculated in cryoSPARC for particle projections is shown. The heat map shows the number of particles for each viewing angle.

**Figure S3.**
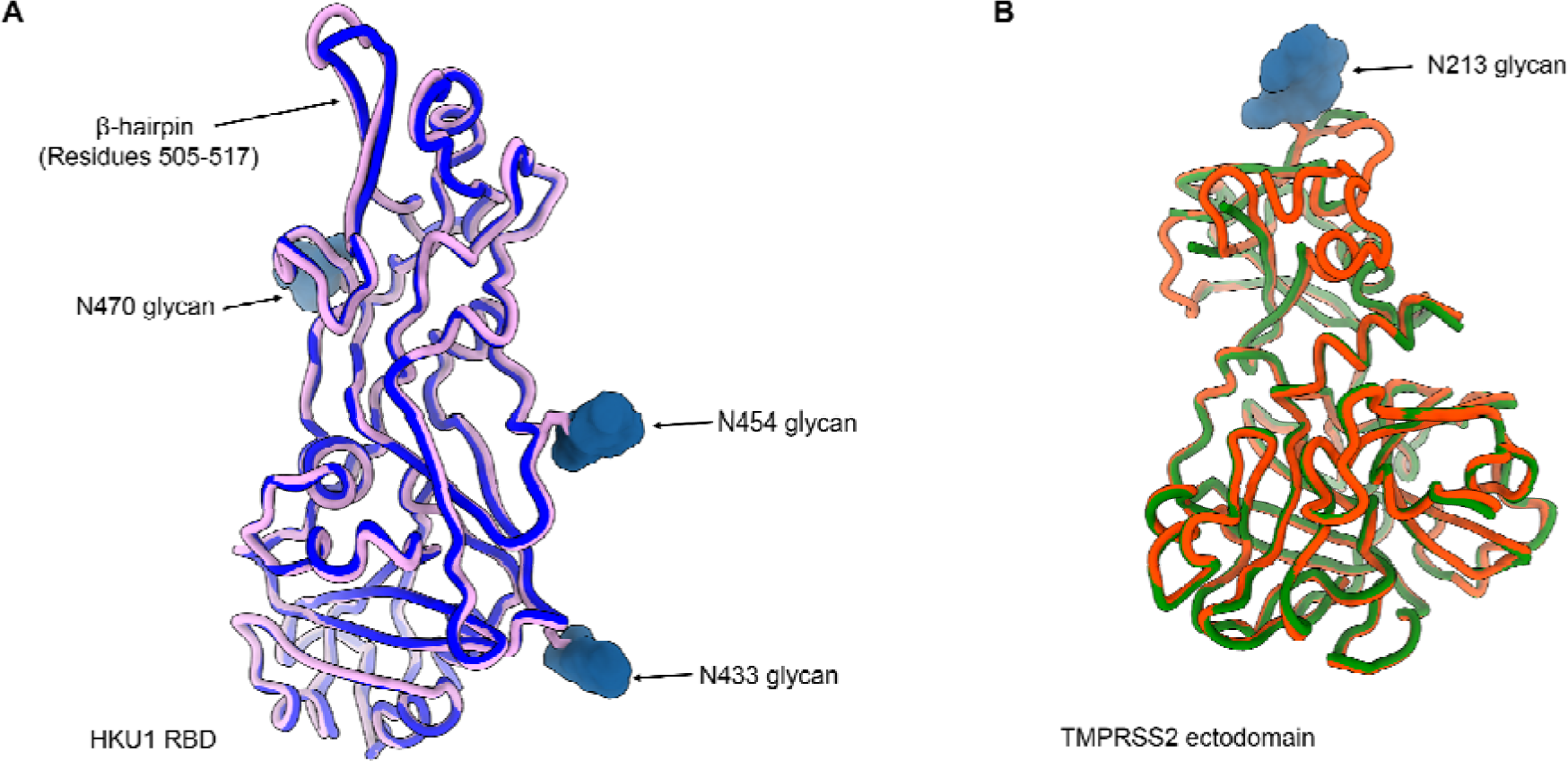
Comparison of the HKU1 RBD and TMPRSS2 structures to the TMPRSS2-bound HKU1 RBD cryoEM structure. (A) Ribbon diagram of the cryoEM structure of the HKU1 RBD (purple) bound to the human TMPRSS2 ectodomain superimposed to the crystal structure of the apo HKU1 RBD (blue, PDB 5KWB). The TMPRSS2 ectodomain is omitted for clarity. (B) Ribbon diagram of the cryoEM structure of the human TMPRSS2 ectodomain (red) bound to the HKU1 RBD superimposed to the crystal structure of the nafamostat-bound TMPRSS2 (green, PDB 7MEQ). The HKU1 RBD is omitted for clarity.

**Figure S4.**
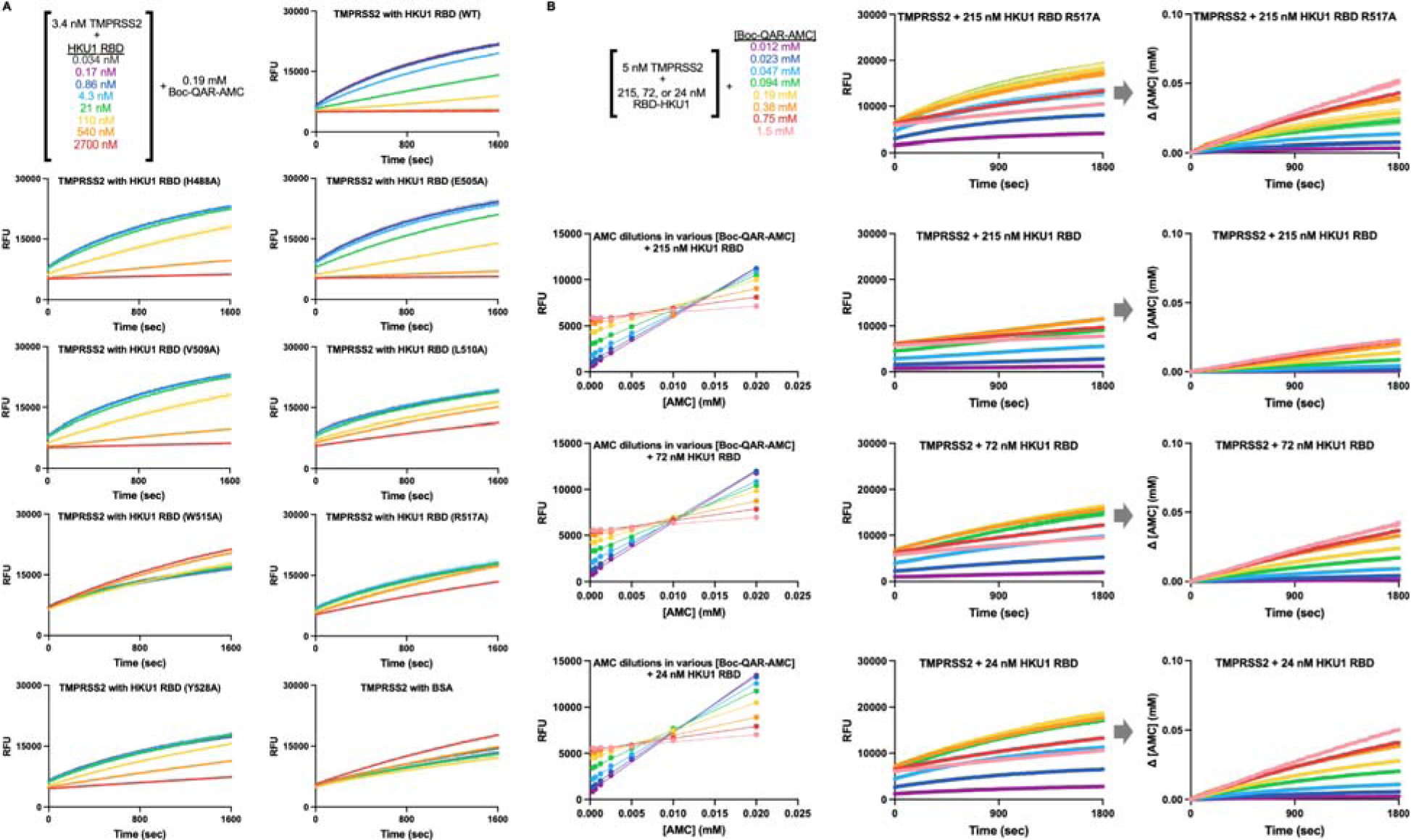
Enzymatic characterization of TMPRSS2 inhibition by the wildtype isolate N1 and alanine interface mutant HKU1 RBDs. (A) Raw data used for Figure 3B; the upper left panel shows the color key used for the other panels. Data show measured fluorescence (RFU) over time with 3.4 nM TMPRSS2 (harboring the N-terminal SUMO fusion, the C379-T447C disulfide bond and the N249 glycan) incubated with 0.19 mM Boc-QAR-AMC at 22°C in the presence of various concentrations of HKU1 isolate N1 RBD and alanine interface mutant RBDs. (B) Raw data and calculations used for Figure 3C-D; the upper left panel shows the color key used for the other panels. Lower left panels show a summary of AMC standard curves used to calculate AMC release from the Boc-QAR-AMC peptide substrate with 215, 72, and 24 nM HKU1 RBD, accounting for the inner filter effect. The middle panels show measured fluorescence (RFU) over time with 5 nM TMPRSS2 and 215, 72, or 24 nM WT HKU1 RBD, or 215 nM HKU1 R517A RBD, incubated with various concentrations of Boc-QAR-AMC at 22 °C. The right panel shows the change in calculated AMC concentration over time.

**Figure S5.**
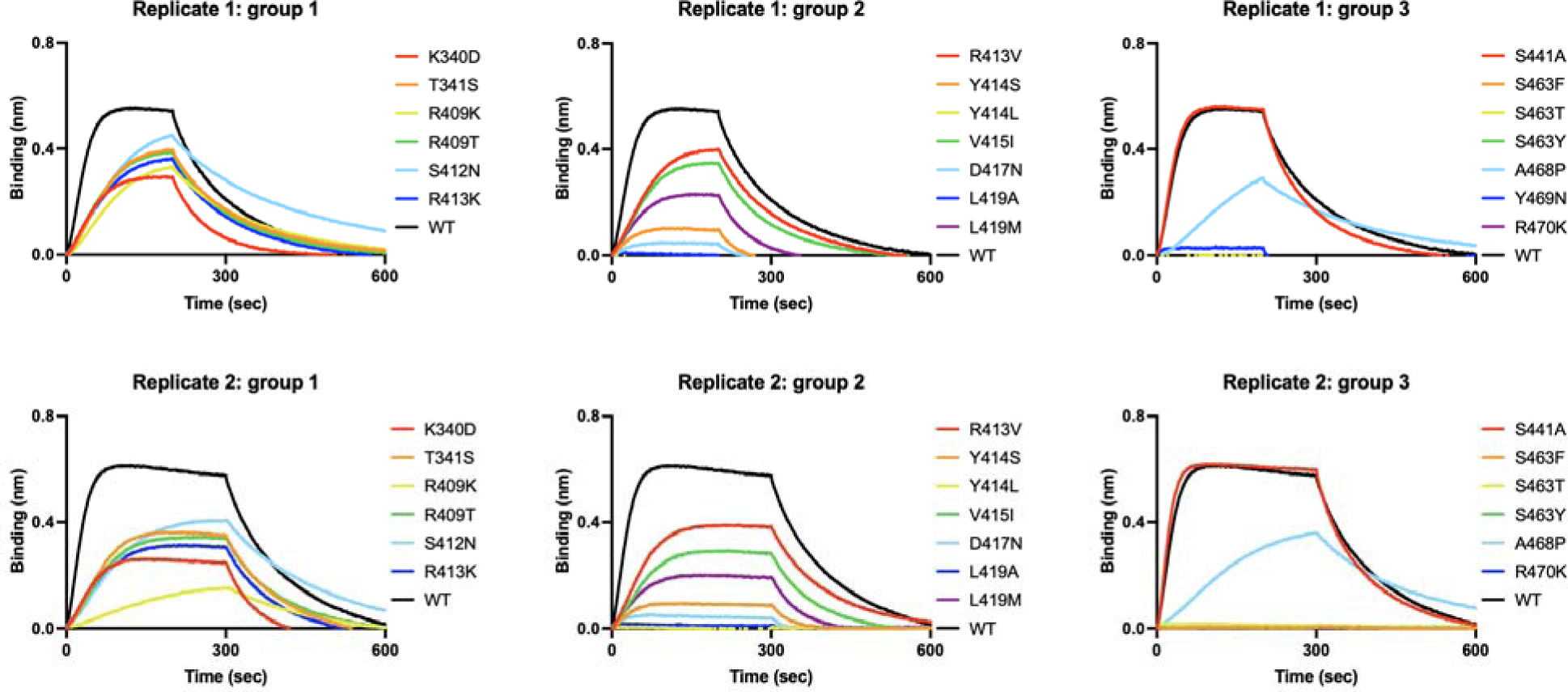
Binding of TMPRSS2 point mutants to the biotinylated HKU1 RBD immobilized on biolayer interferometry SA biosensors. The top and bottom rows show baseline-subtracted response curves for the first and second biological replicates. For the first replicate, biotinylated HKU1 RBD-loaded SA tips were dipped into 100 nM TMPRSS2 for 200 seconds followed by dissociation for 500 seconds. For the second replicate, biotinylated HKU1 isolate N1 RBD-loaded SA tips were dipped into 100 nM TMPRSS2 for 300 seconds followed by dissociation for 500 seconds. To facilitate visualization, the binding curves from point mutants are split into three groups and include wildtype (WT) TMPRSS2 for reference.

**Figure S6.**
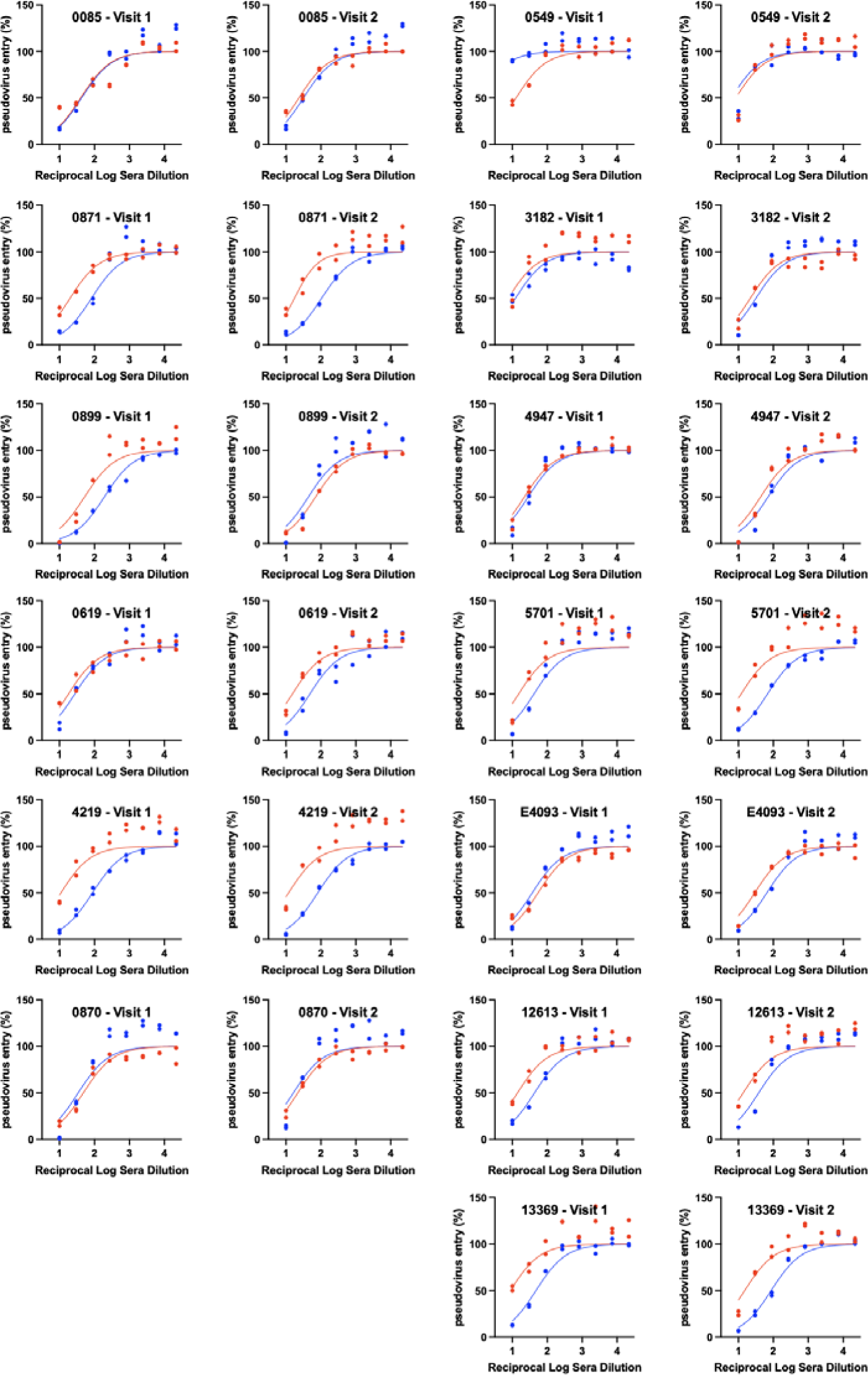
Evaluation of HKU1 infection-elicited serum neutralizing activity in humans. Dose-response curves for neutralization of VSV pseudotyped with wildtype isolate N1 HKU1 S (WT, red) or the N355 glycan knockout (S357A, blue) mutant by human serum antibodies, shown in red and blue, respectively. Visits 1 and 2 correspond to blood draws at the time of PCR-positive testing and 30 days later, respectively^52, 53^.

**Table S1.**
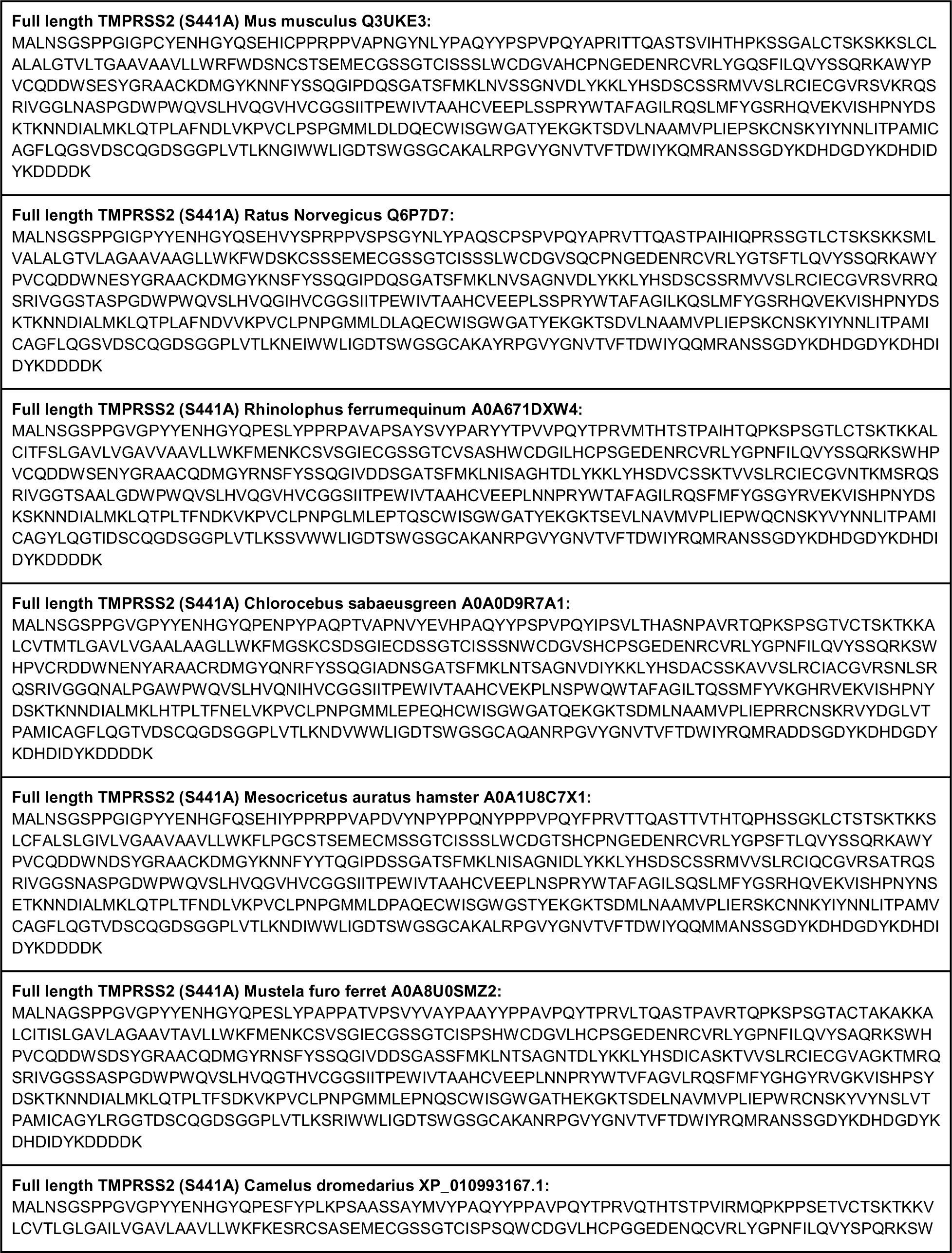
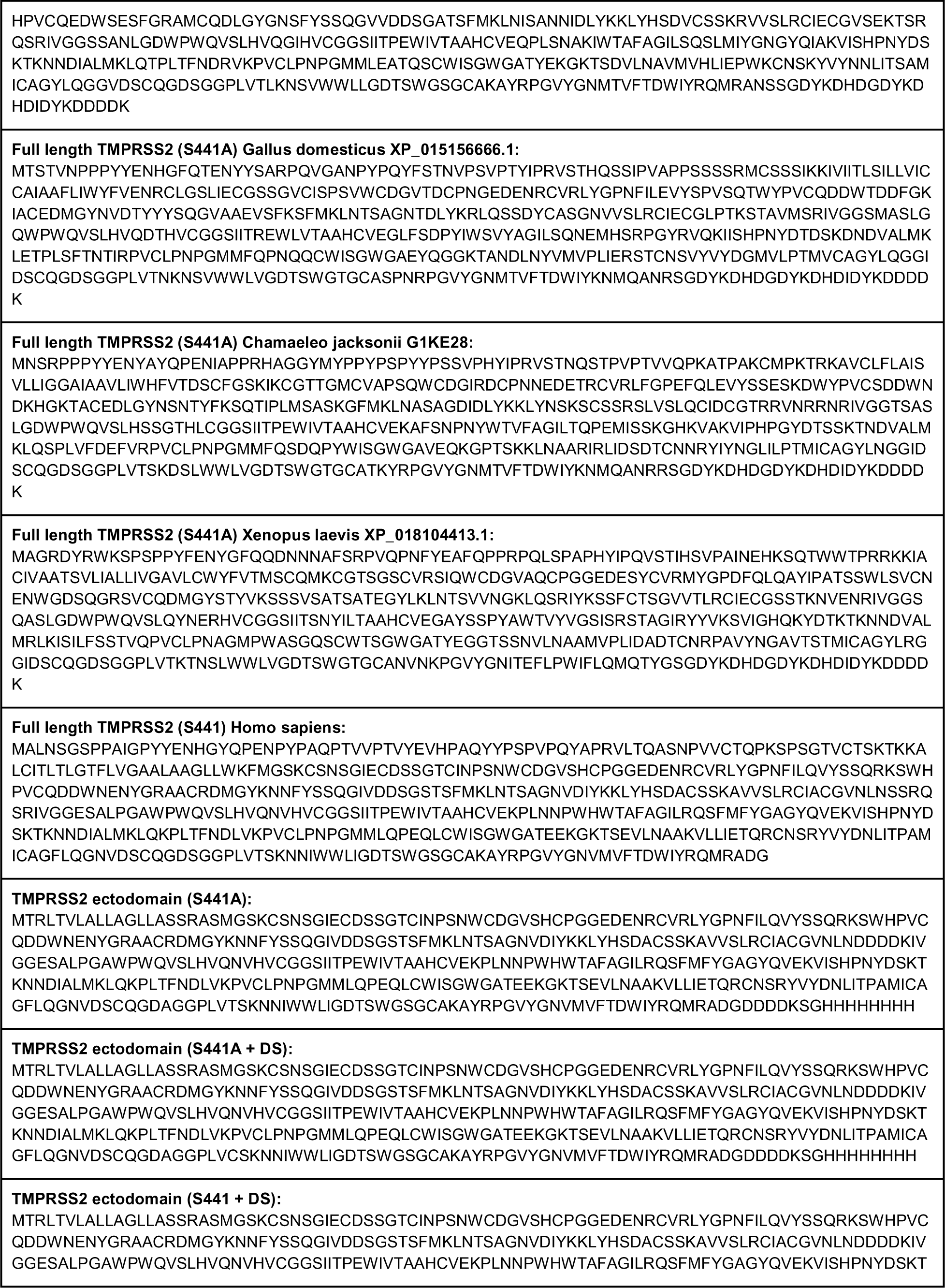
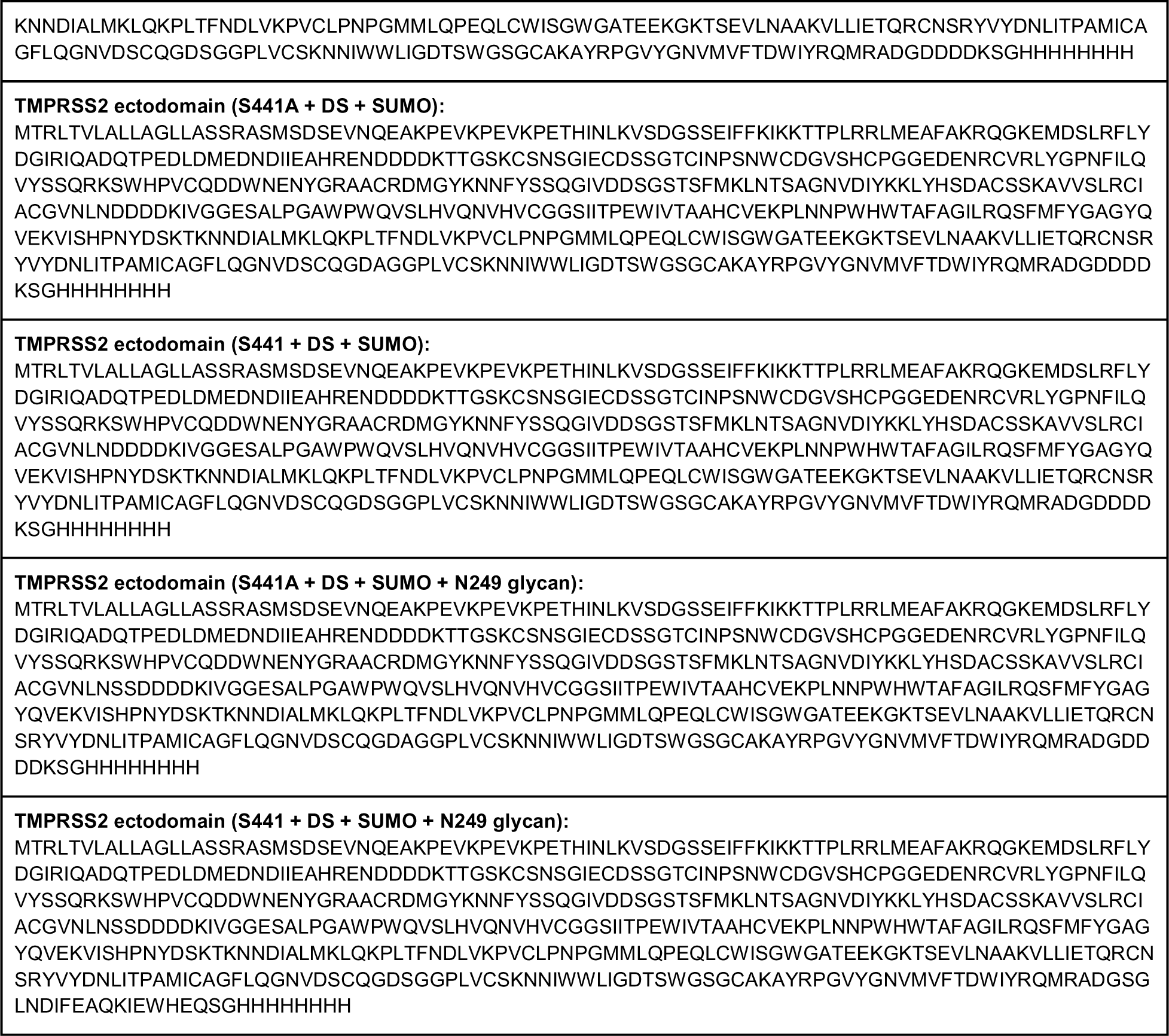
Amino acid sequences of designed TMPRSS2 constructs.

**Table S2.**
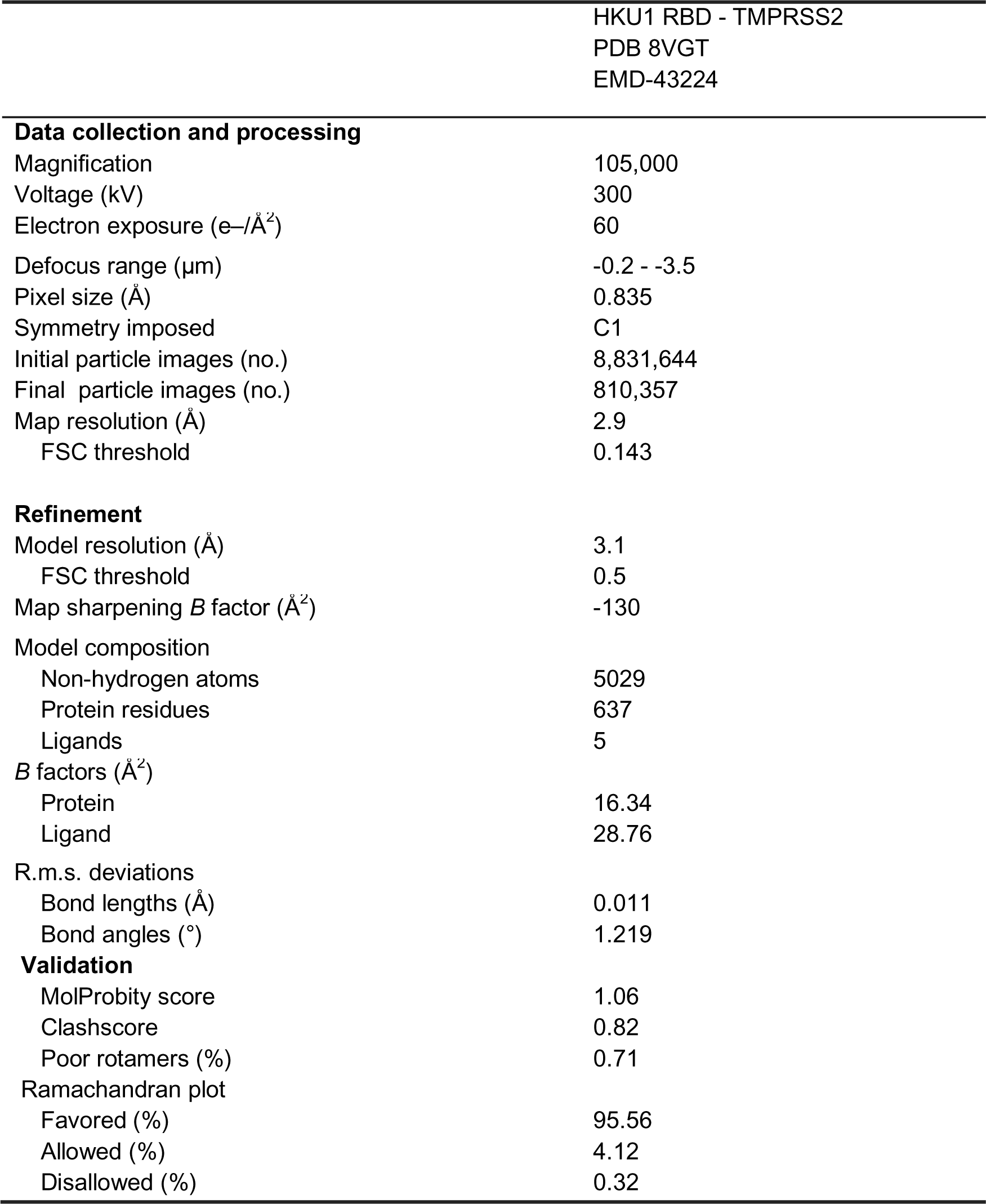
Cryo-EM data collection, refinement and validation statistics.

|  |  |
| --- | --- |
|  | HKU1 RBD - TMPRSS2<br>PDB 8VGT<br>EMD-43224 |
| **Data collection and processing** |  |
| Magnification | 105,000 |
| Voltage (kV) | 300 |
| Electron exposure (e<sup>-</sup>/Å<sup>2</sup>) | 60 |
| Defocus range (μm) | -0.2 - -3.5 |
| Pixel size (Å) | 0.835 |
| Symmetry imposed | C1 |
| Initial particle images (no.) | 8,831,644 |
| Final particle images (no.) | 810,357 |
| Map resolution (Å) | 2.9 |
| FSC threshold | 0.143 |
| **Refinement** |  |
| Model resolution (Å) | 3.1 |
| FSC threshold | 0.5 |
| Map sharpening *B* factor (Å<sup>2</sup>) | -130 |
| Model composition |  |
| Non-hydrogen atoms | 5029 |
| Protein residues | 637 |
| Ligands | 5 |
| *B* factors (Å<sup>2</sup>) |  |
| Protein | 16.34 |
| Ligand | 28.76 |
| R.m.s. deviations |  |
| Bond lengths (Å) | 0.011 |
| Bond angles (°) | 1.219 |
| **Validation** |  |
| MolProbity score | 1.06 |
| Clashscore | 0.82 |
| Poor rotamers (%) | 0.71 |
| Ramachandran plot |  |
| Favored (%) | 95.56 |
| Allowed (%) | 4.12 |
| Disallowed (%) | 0.32 |

**Data S1. Human TMPRSS2 SNPs at the HKU1 RBD-interacting site present in the GnomADv4 database.**

**Data S2. Human TMPRSS2 SNPs at the HKU1 RBD-interacting site present in the Regeneron Million Exome Variant database.**

**Data S3.** Demographic data for HKU1 infected serum donors.

| Manuscript code | Country of isolation | Age | Gender | Collection date V1 | Collection date V2 |
| --- | --- | --- | --- | --- | --- |
| HK-01 | Netherlands | 47 | 2 | 2/3/10 | 3/5/10 |
| HK-02 | Spain | 36 | 2 | 3/16/10 | 4/20/10 |
| HK-03 | Spain | 27 | 2 | 3/5/10 | 4/26/10 |
| HK-04 | Spain | 38 | 2 | 4/8/10 | 5/12/10 |
| HK-05 | Poland | 56 | 1 | 3/8/10 | 4/16/10 |
| HK-06 | Poland | 54 | 2 | 1/21/10 | 2/19/10 |
| HK-07 | Poland | 48 | 1 | 3/3/10 | 4/2/10 |
| HK-08 | Poland | 28 | 2 | 2/23/10 | 4/19/10 |
| HK-09 | Poland | 35 | 1 | 3/4/10 | 4/26/10 |
| HK-10 | Poland | 47 | 1 | 2/10/10 | 3/15/10 |
| HK-11 | Poland | 26 | 2 | 2/8/10 | 3/11/10 |
| HK-12 | United Kingdom | 53 | 2 | 12/11/07 | 1/18/08 |
| HK-13 | Spain | 27 | 2 | 2/16/09 | 4/16/09 |

